# Deciphering interferon functions in avian influenza: Insights from receptor knockout models in the natural host

**DOI:** 10.1101/2025.01.24.634794

**Authors:** Mohanned Naif Alhussien, Hanna-Kaisa Vikkula, Romina Klinger, Christian Zenner, Simon P Früh, Rashi Negi, Theresa von Heyl, Sabrina Schleibinger, Milena Brunner, Tom VL Berghof, Leora Avolio, Arne Reich, Benjamin Schade, Bassel A. Abukhadra, Silke Rautenschlein, Rudolf Preisinger, Hicham Sid, Benjamin Schusser

## Abstract

The rapid cross-species transmission of highly pathogenic avian influenza presents a significant zoonotic threat. Elucidating the avian interferon (IFN) system, the primary antiviral defense in chickens, is critical for controlling the virus at its source and preventing its spillover into humans and other species. We engineered type I (IFN-α/β) and type III (IFN-λ) IFN receptor knockout chickens to dissect the role of IFNs in viral infections. Results revealed that type I IFN predominantly modulates innate immune cell populations, T cell subsets, and their contribution to antibody production following immunization under physiological conditions. *In ovo* and *in vivo* challenge experiments utilizing diverse influenza A virus strains demonstrated strain-specific roles of both IFN-α/β and IFN-λ in orchestrating viral pathogenesis, immunological responses, and tissue-tropism effects. Notably, type I IFN was particularly crucial in the initial defense mechanisms against H3N1 avian influenza A virus infection. These novel models offer unprecedented insights into avian IFN biology within the context of avian influenza, which is essential for developing more effective strategies to prevent and control this public health challenge.

## Introduction

Chickens are highly susceptible to viral pathogens such as avian influenza A virus (AIV), a zoonotic virus transmissible to humans, which causes significant economic losses in the poultry industry^1, 2, 3^. In 2022, 67 countries across five continents reported outbreaks of H5N1 highly pathogenic avian influenza, losing over 131 million domestic poultry. The disease continued to spread as 14 more countries reported outbreaks, primarily across the Americas^4^. The emergence of various zoonotic avian influenza strains has heightened global concerns about the potential for these strains to mutate and sustain human-to-human transmission, possibly leading to a pandemic^5, 6^. Despite substantial advancements in our comprehension of the mammalian host response to pathogenic insults, knowledge of immune mechanisms in avian species, the natural hosts of the virus, remains limited. As a result, strategies to control AIV have heavily relied on biosecurity measures and depopulation of infected flocks, imposing substantial economic burdens on the poultry industry and amounting to billions of dollars in losses^7^. The insufficiency of these traditional approaches in managing the growing prevalence of highly pathogenic AIV outbreaks has driven interest in alternative antiviral therapies^2, 3^. Notably, France and the Netherlands have implemented domestic duck vaccination as a control measure against highly pathogenic avian influenza outbreaks^8^.

Interferons (IFNs) were first identified in chicken embryos. These proteins are characterized by their ability to inhibit viral replication, a property that led to their designation^9^. IFNs are cytokines that induce the expression of a myriad of IFN-stimulated genes (ISGs) that act on innate and adaptive immunity, eliciting an antiviral state against invading viral pathogens^10, 11^. They are classified based on their molecular structure, receptor specificity, and induction pathways. In chickens, type I IFNs (IFN-α and IFN-β) and type III IFN (IFN-λ) play critical roles, with IFN-α and IFN-λ being the primary focus of immunological studies due to their significant involvement in antiviral responses. To accomplish the primary function of IFNs, these cytokines need to bind to their respective receptors. The type III IFN receptor is a heterodimer consisting of the ubiquitously expressed receptor chain IL-10Rβ and the epithelium-specific receptor chain IFNLR1 (IL-28Rα). In contrast, type I IFN (IFNAR1/IFNAR2) receptors demonstrate widespread expression across diverse tissues and cell types throughout the body^12, 13^.

The molecular cloning and characterization of chicken IFN-λ were relatively recent, uncovering antiviral properties comparable to those of mammalian IFN-λ^11, 14^. However, despite this progress, the mechanisms and functional roles of type I and III IFNs in chickens remain largely uncharacterized. Murine IFN-λ is the initial IFN produced during a viral invasion, providing epithelial antiviral protection against low viral loads without eliciting an inflammatory response^15^. When the infection surpasses the control of IFN-λ, type I IFN is activated, associated with an inflammatory response and immunopathology. Unlike the widespread effects of type I IFN, the similarity of IFNLR1 expression with the tissue tropism of influenza A virus allows mammalian IFN-λ to specifically target cell types at risk of infection. This effectively induces antiviral genes in these cells, aiding in controlling influenza A virus spread without overstimulating the immune system and causing immunopathology^16, 17^. IFNAR1 knockout (KO) mice show increased susceptibility to several viral infections, but not influenza. However, mice lacking IFNAR1 and IFNLR1 cannot control influenza infection^18, 19^. Researchers have shown that mammalian IFN-λ can also have pro-inflammatory effects by disrupting the lung epithelial barrier, hindering lung repair, and impairing epithelial proliferation during influenza infections^20, 21^.

Despite the discovery of IFNs in chickens, our knowledge of the avian IFN system remains limited^9^. Unraveling the complexities of the IFN system is crucial for devising strategies to control avian influenza in its natural host, thereby preventing zoonotic transmission to humans and other species. This limitation is primarily due to the complex and overlapping signaling pathways of type I and type III IFNs, which induce similar sets of ISGs and produce almost indistinguishable biological activities despite signaling through distinct receptors^10^. Furthermore, our understanding of ISGs and their antiviral functions in chickens is primarily based on research from humans and other mammals, which may not fully account for the fundamental differences in innate immunity between chickens and mammals^11, 13, 22^. The lack of appropriate biotechnological tools, such as IFN receptors deficient genetically engineered chickens, hinders progress in this field. These challenges make it difficult to elucidate the individual antiviral and immunomodulatory roles of type I and type III IFNs in avian research despite the potential value of these cytokines as targets for developing therapeutics against a broader range of viruses^13, 23^.

To address this gap, we utilized chicken primordial germ cells (PGCs) to generate IFNAR1 and IFNLR1 knockout (KO) chicken lines. These genetically modified lines allow for the selective study of type I and type III IFN responses. Through experiments conducted *in vitro*, *in ovo*, and *in vivo*, we investigated the impact of these IFN classes on various physiological functions, antibody responses to immunization, and host-pathogen interactions. Notably, *in vivo* H3N1 infections revealed distinct and novel mechanisms for both types of chicken IFN in the context of H3N1 defense. These newly developed chicken lines provide critical insights into the roles of IFN-α/β and IFN-λ in immune responses during various viral infections, including influenza and infectious bronchitis virus (IBV). The knowledge gained from these studies is fundamental for developing therapeutic methods and immunomodulatory strategies to control and eradicate the significant public health challenges in avian species.

## Results

### Successful generation of IFN receptor KO chickens

Knocking out the IFNAR1 or IFNLR1 receptors generates animal models deficient in type I or type III IFN signaling, respectively. This receptor-targeted approach, in contrast to the cytokine-targeted approach, is especially effective in chickens, where multiple IFN-α genes exist, as it blocks the entire signaling pathway, allowing for comprehensive studies of IFN’s roles in immunity and viral responses^12, 13, 24^. Chickens lacking either the receptor of IFN-α/β (IFNAR1) or IFN-λ (IFNLR1) were generated by designing guide RNAs to target the coding region within exon 1 of the IFNAR1 chain of the IFN-α/β receptor and the epithelium-specific chain (IFNLR1) of the IFN-λ receptor. PGCs, precursors of sperms and oocytes, were co-transfected with a CRISPR/Cas9 vector expressing a single guide RNA and Cas9-2A-eGFP targeting the coding region within exon 1 of the respective receptor genes (Fig. 1a, b). To generate the IFNAR1 knockout, homology-directed repair (HDR) utilizing single-stranded oligodeoxynucleotide (ssODN) templates was used to facilitate the targeted gene edits. Transient expression of eGFP in transfected PGCs allowed for the selection of successfully transfected cells via automated cell sorting. Subsequent Sanger sequencing confirmed several clonal PGCs lines with the desired modifications for IFNAR1 and IFNLR1.

**Fig. 1.**
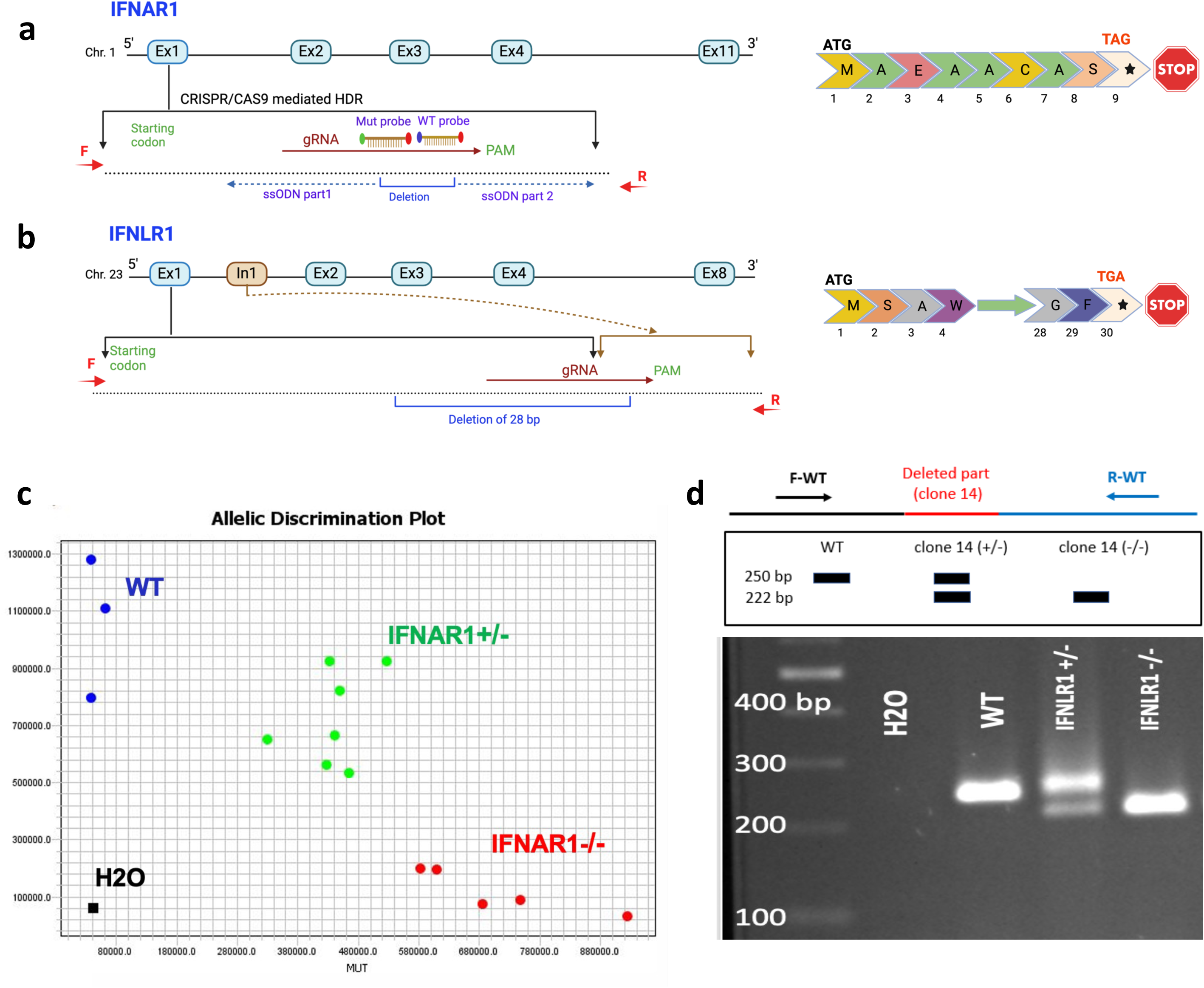
Targeting strategy and genotype for knockout of chicken IFN-α/β receptor (IFNAR1) and IFN-λ receptor (IFNLR1). **a**, Guide RNAs designed to target the coding region within exon 1 of the IFNAR1 chain of the IFN-α/β receptor and the epithelium-specific chain (IFNLR1) of the IFN-λ receptor. For IFNAR1, the INDEL was pre-designed using CRISPR-mediated HDR technology. The IFNAR1 KO clone has 7 bp deletion and produces a stop codon after 21 bp. **b,** For IFNLR1 positive PGCs clones, the clone with the maximum deletion (28 bp) was selected as it is suitable to design IFNLR1 KO specific PCR genotyping assay and produce a stop codon after 87 bp. **c**, TaqMan genotyping assay for IFNAR1. **d,** PCR genotyping assay for IFNLR1.

For IFNLR1, the PGC clone with the largest deletion (28 bp) was selected, leading to a stop codon formation after 87 bp. Similarly, for IFNAR1, the clone with a 7 bp deletion produced a stop codon after 21 bp (Fig. 1a, b). These clones were used for the generation of germline chimeras. Selected chimeric roosters demonstrated 4% and 7% germline transmission rates for IFNAR1 and IFNLR1 modifications, respectively, resulting in heterozygous IFNAR1^+/−^ and IFNLR1^+/−^ progeny. The modified alleles segregated in a Mendelian manner, and homozygous IFNAR1^−/−^ and IFNLR1^−/−^ lines were successfully established. TaqMan allelic discrimination assays were employed to genotype the IFNAR1 KO animals (Fig. 1c), while a PCR genotyping assay was established for IFNLR1 KO chickens (Fig. 1d). Phenotypically, both IFNAR1^−/−^ and IFNLR1^−/−^ chickens exhibited normal body weight gain compared to their WT counterparts (Fig. 2a). Since both type I IFN and type III IFN play crucial roles in protecting the female reproductive tract against viral infections and maintaining immune homeostasis in mammals^25, 26^, we evaluated the fertility of IFNAR1^−/−^ and IFNLR1^−/−^ chickens by breeding homozygous birds from each line separately. The resulting F2 generation revealed normal egg production and fertility in healthy, unchallenged KO chickens (data not shown), consistent with studies in mice lacking either IFNAR1^−/−^ or IFNLR1^−/−27, 28^.

**Fig. 2.**
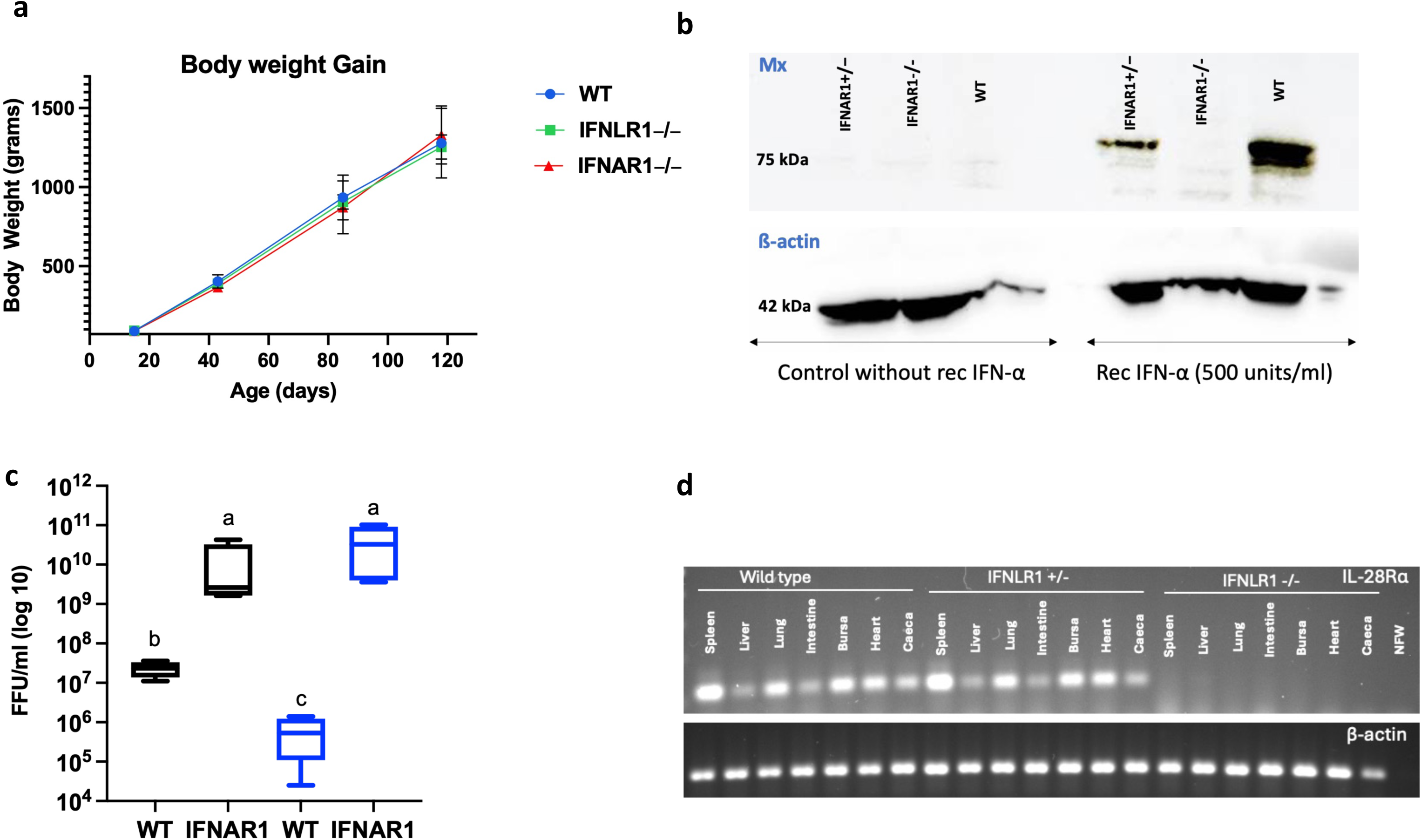
Growth, and successful generation of genetically modified chickens. a, Body weight development of WT, IFNLR1^−/−^ and IFNAR1^−/−^ chickens. **b,** Western blot confirms successful knockout of IFNAR1: WT, IFNAR1^+/−^, and IFNAR1^−/−^ chicken embryonic fibroblast cells (n = 3) were cultured in a 6-well plate and stimulated with recombinant IFN-α (500 U/ml) for 12 hours. For the control groups, CEF cells were treated with CEF medium without recombinant IFN-α. After stimulation, the control and treatment groups’ CEF cells were trypsinized and collected for an SDS-Gel and Western Blot experiment. The results demonstrate functional loss of type I IFN receptor (IFNAR1) in CEF IFNAR1^−/−^ cells treated with recombinant IFN-α, as the Mx protein (75 kDa) was observed solely in the CEF WT and IFNAR1^+/−^ cells treated with recombinant IFN-α. **c,** WSN33 viral titration in WT and IFNAR1^−/−^ embryo pretreated with recombinant IFN-α. ED10 chicken embryos (blue columns) were stimulated with 1.5 × 10^5^ U of recombinant IFN-α in 100 μl PBS 12 h before and at the time of infection. As controls (black columns) did not receive IFN-α stimulation. All the groups of embryos were infected with 1000 FFU of WSN33, and viral load in the allantoic fluid was analyzed 24 h post-infection by titration on MDCK cells. Number of embryos, for the control groups (WT = 4, IFNAR1^−/−^ = 4), and treatment groups (WT = 4, IFNAR1^−/−^ = 4). Significance was calculated by one-way ANOVA. Statistically different groups (P ≤ 0.05) are indicated with different superscript letters (a, b, c). **d,** RT-PCR confirms functional loss of IFNLR1 in homozygous embryos: expression analysis of IFN-λ receptor (IL-28Rα) was studied in different tissues of 18-day-old embryos challenged with H3N1 to ensure induced expression of IL-28Rα. RNA was isolated via the Trizol method after the genotyping of embryos. cDNA was transcribed via GoScript Transcription Mix, Random Primers. PCR was performed with β-actin primers as control and IL-28Rα primers to check the presence of IL-28Rα receptors in tissues. Nuclease-free water (NFW) was used as a negative control for both. A 1.5% tris-borate-EDTA (TBE) was used to visualize β-actin amplicon at 300 bp and 2% TBE to visualize 108 bp for IL-28Rα.

### Knockout of IFNAR1 and IFNLR1 impairs IFN signaling and induction of antiviral state

Western blot analysis confirmed the functional loss of the IFNAR1 receptor in the IFNAR1^−/−^ chickens. Treatment with recombinant IFN-α resulted in the induction of IFN-stimulated protein Mx in WT and IFNAR1^+/−^ chicken embryo fibroblasts (CEF) but not in the CEF derived from IFNAR1^−/−^ embryos (Fig. 2b). In addition, treatment of both WT and IFNAR1^−/−^ embryos with recombinant IFN-α before WSN33 (H1N1) challenge resulted in a significant (P ≤ 0.05) reduction of viral titers in the allantoic fluid of WT but not in IFNAR1^−/−^ embryos (Fig. 2c). These results demonstrated that the signaling pathway mediated by IFNAR1 is non-functional in the absence of this receptor, thus validating the knockout model. Additionally, RT-PCR analysis was conducted to assess the expression of IFNLR1 (IL-28Rα) across multiple tissues, including the spleen, liver, lung, intestine, bursa of Fabricius, heart, and cecum in WT, IFNLR1^+/−^, and IFNLR1^−/−^ embryos. The analysis revealed a complete absence of IFNLR1 mRNA in all IFNLR1^−/−^ tissues compared to detectable levels in WT and IFNLR1^+/−^ samples. This absence indicates the successful functional loss of IFNLR1 (Fig. 2d).

#### Knockout of IFNAR1 and IFNLR1 modulates the population of immune cells in the blood and spleen

Flow cytometry analysis demonstrated a significant (P ≤ 0.05) reduction in the blood B cells and monocyte populations and an increase in αβ T cells in one-month-old IFNAR1^−/−^ chickens compared to WT and IFNLR1^−/−^. Furthermore, there was a marked decrease in CD8^+^ γδ T cells, accompanied by a significant (P ≤ 0.05) increase in CD4^+^ αβ T cells in the IFNAR1^−/−^ group compared to other groups (Fig. 3, Supplementary Fig. 1). Although flow cytometry analysis did not reveal significant differences in the number of B cells, macrophages, αβ, and γδ T cells in the spleen of WT, IFNAR1^−/−^ and IFNLR1^−/−^, there was a considerable decrease in CD8^+^ γδ T cells in IFNLR1^−/−^ as compared to WT chicks (Fig. 3). These findings highlight the differential impact of IFNAR1 and IFNLR1 KO on specific T cell subsets and other immune cell populations, and the important roles of these receptors in immune regulation.

**Fig. 3.**
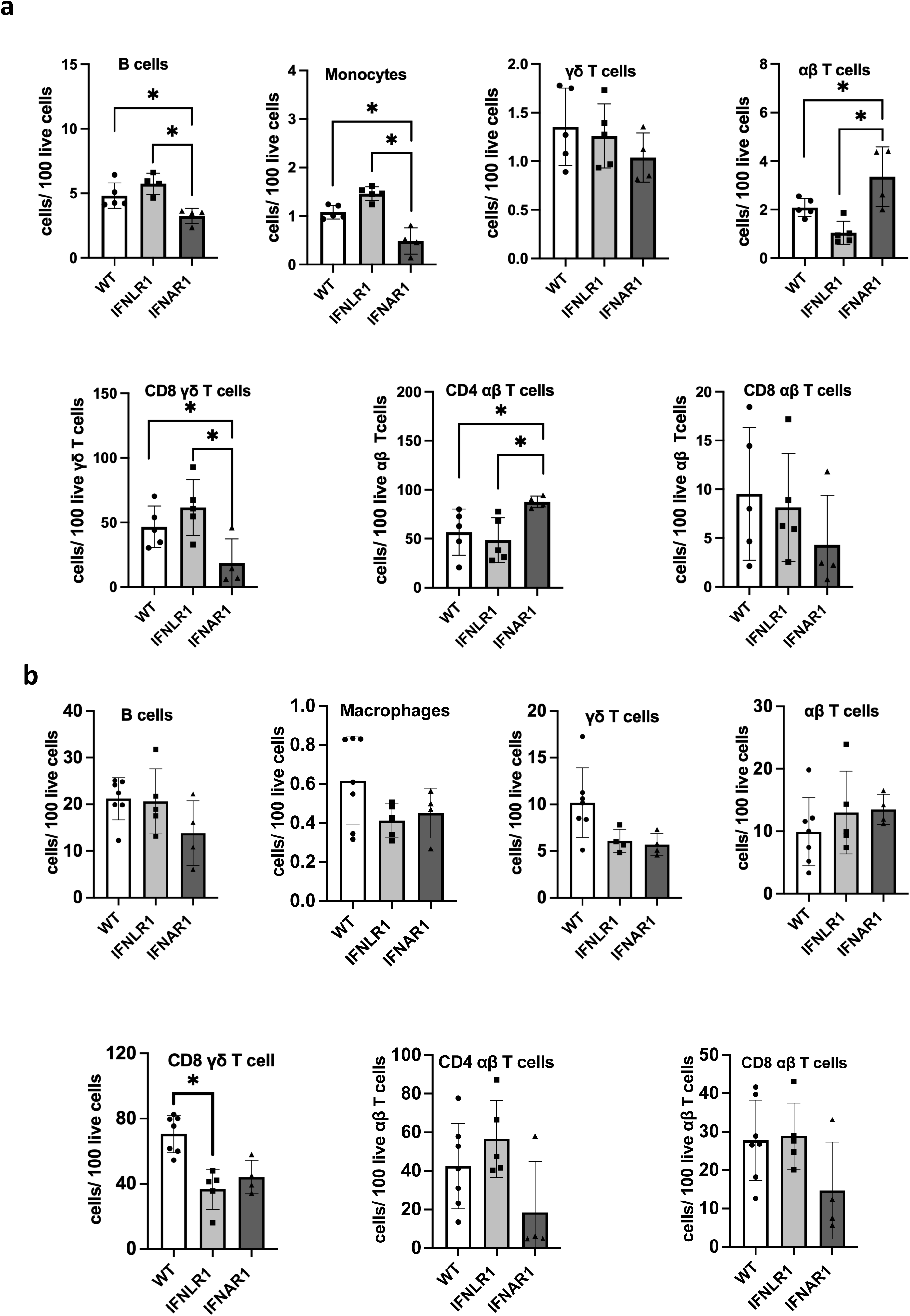
Flow cytometry analysis of PBMCs and splenocytes of WT, IFNLR1^−/−^ and IFNAR1^−/−^ in one-month-old chickens. a, PBMCs and **b,** splenocytes were isolated and analyzed for differential immune cell populations, including monocytes/macrophages, B cells, αβ TCR2,3+ or γδ TCR1 + T cells, and associated CD4 and CD8 T cells subpopulations. One-way ANOVA was used to analyze the difference between the groups (**\*,** P ≤ 0.05). Number of animals for the analysis of PBMCs (WT = 5, IFNLR1^−/−^ = 5, IFNAR1^−/−^ = 4), and analysis of splenocytes (WT = 7, IFNLR1^−/−^ = 5, IFNAR1^−/−^ = 4).

#### Knockout of IFNLR1 did not modulate the cecum microbiome

Given the high expression of IFNLR1 in epithelium-rich organs such as the intestine and considering the constant microbial exposure in this region, we investigated the impact of IFNLR1 knockout on the cecum microbiomes of one-month-old healthy chicks. While microbial sequencing disclosed distinct multidimensional scaling plots between the IFNLR1^−/−^ and their WT counterparts, subsequent alpha diversity analyses, and taxonomic diversity, did not demonstrate significant differences between the groups (Supplementary Fig. 2). However, it is noteworthy that this difference could be more pronounced during immune challenges when IFN-λ signaling is actively engaged.

### Reduced IgM and IgY production in IFNAR1^−/−^ and IFNLR1^−/−^ chickens following immunization

To study the role of type I and type III IFNs in immune response and antibody production, one-month-old chicks were immunized with Keyhole Limpet Hemocyanin (KLH) (Supplementary Fig. 3). Analysis of total IgM and IgY concentrations in the plasma of preimmunized animals (day 0) revealed a significantly (P ≤ 0.05) reduced IgY synthesis and secretion in IFNAR1^−/−^ chickens compared to the WT group. Furthermore, all groups showed significant (P ≤ 0.05) increases in total IgM and IgY concentrations on day 5 after both primary and secondary immunizations compared to day 0. These elevations were solely attributed to immunization with KLH and FIA (Fig. 4a, b). Analysis of KLH-specific IgM and IgY levels showed significantly (P ≤ 0.05) lower levels of both antibodies in IFNAR1^−/−^ and IFNLR1^−/−^ chicks compared to the WT group. Notably, although IFNLR1^−/−^ chicks had significantly (P ≤ 0.05) lower IgM and IgY levels than IFNAR1^−/−^ on day 5 post-primary immunization, they were able to reach immunoglobulin levels comparable to WT chicks by day 5 after booster immunization (Fig. 4c, d). This indicates that IFN-λ contributes to the initial antibody production following immunization but can be compensated for by type I IFN during booster immunization.

**Fig. 4.**
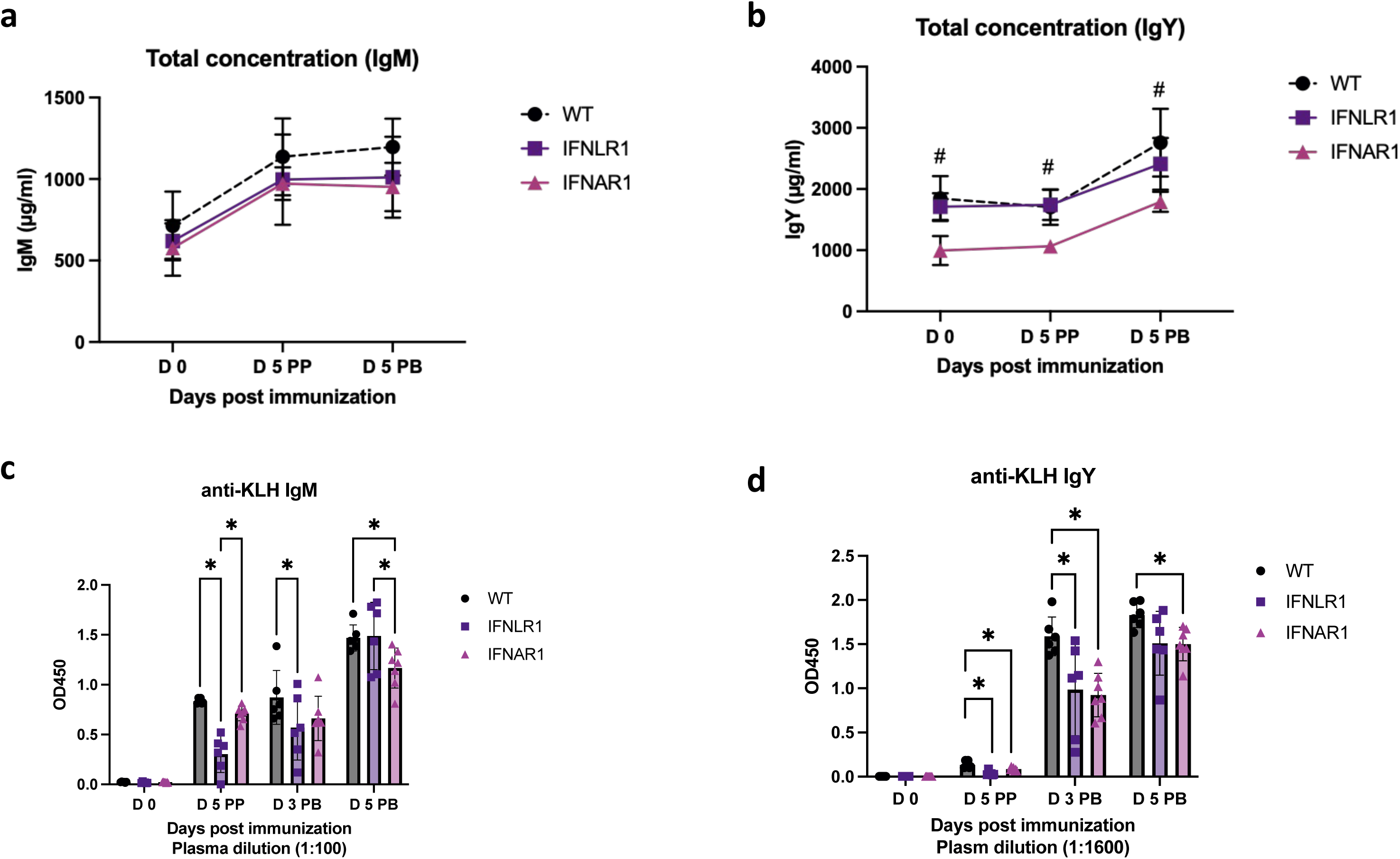
Reduced IFNLR1^−/−^ and IFNAR1^−/−^ chicken ability to produce IgM and IgY in response to immunization. Five-week-old chickens were immunized via intramuscular injection with 300 µg KLH, mixed 1:1 with incomplete Freund’s adjuvant. A booster injection of 300 µg KLH, mixed 1:1 with Freund’s incomplete adjuvant, was administered two weeks after the initial immunization. Blood plasma samples were collected from all groups before immunization (day 0), on day 5 post-primary immunization (PP), and on days 3 and 5 post-booster immunization (PB). **a-d,** Total concentration of IgM, IgY, and KLH antigen-specific IgM and IgY levels were assessed using ELISA. Data are presented as mean and SEM of at least 6 birds per genotype, with the same animals, tracked over time. Statistical differences between groups are indicated by asterisks (**\*,** P ≤ 0.05). # Indicate significant difference (P ≤ 0.05) for IFNAR1^−/−^ vs. WT and IFNLR1^−/−^.

### IFNAR1 knockout reduces MHCII^+^ cells without altering MHCII expression

Analysis of splenocytes isolated on day 5 post-secondary immunization revealed a significant (P ≤ 0.05) reduction in the frequency of MHCII^+^ cells in IFNAR1^−/−^ chickens compared to other genotypes. This decrease was particularly pronounced in specific immune cell subsets, notably MHCII^+^ B cells and MHCII^+^ macrophages, where the percentages of these cells were markedly lower in IFNAR1^−/−^ chickens (Supplementary Figs. 4 and 5). Interestingly, the mean fluorescence intensity (MFI) of MHCII, which indicates the density of MHCII molecules on individual cells, remained consistent across all experimental groups (data not shown). This observation suggests that the observed reduction in MHCII^+^ cells is primarily due to a decrease in the overall populations of macrophages and B cells expressing MHCII rather than a diminished expression of MHCII per cell.

**Fig. 5.**
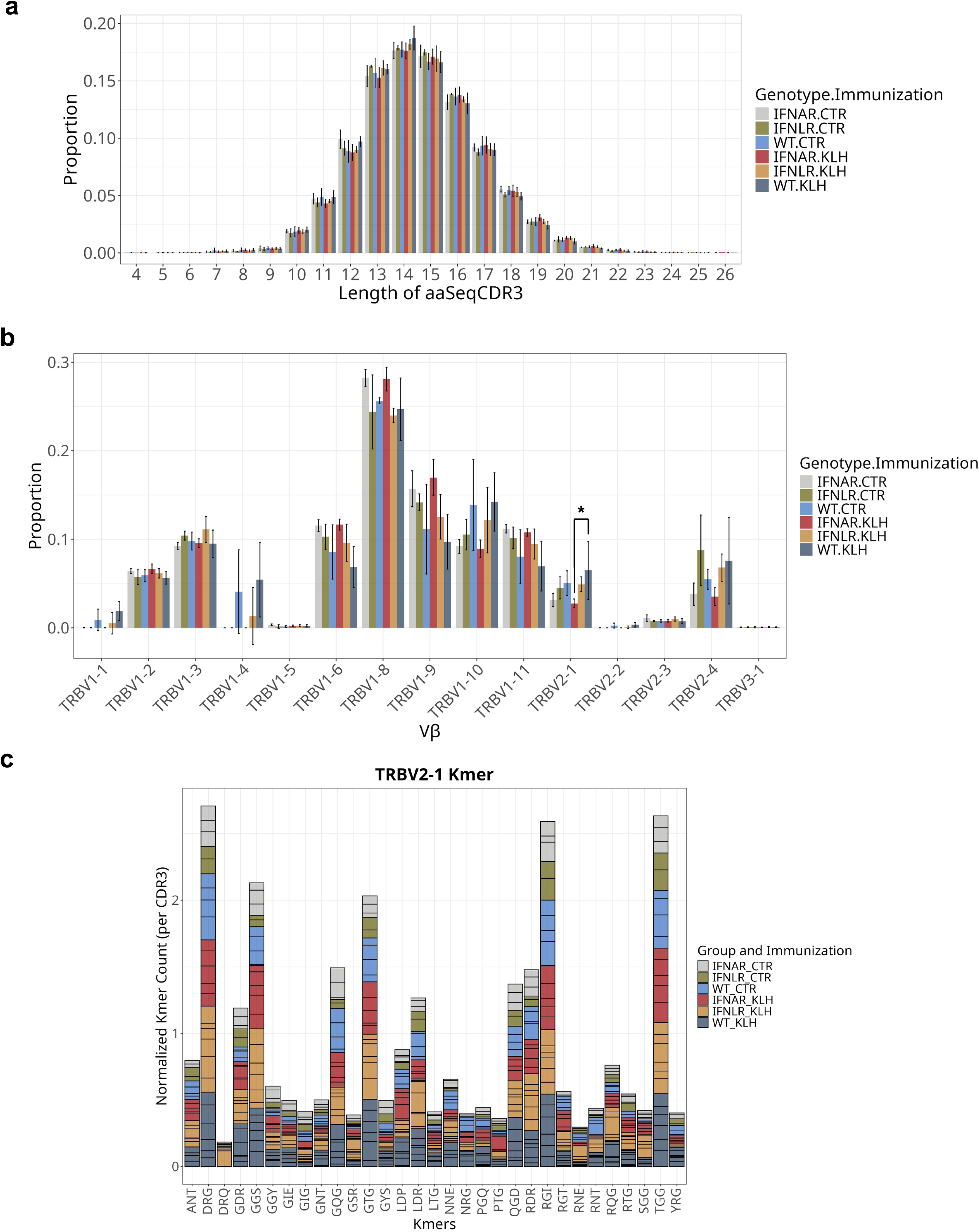
Overall similar splenic TCRβ repertoire across genotypes with reduced TRBV2-1 responses in KLH-immunized IFNAR1^−/−^ chickens. TCRβ CDR3 clonotypes in IFNAR1^−/−^, IFNLR1^−/−,^ and WT animals in the steady state (CTR) or at day 5 post-KLH booster immunization. **a,** CDR3 Spectratype; **b,** Vβ gene usage; **c,** Counts of the 20 most prevalent 3-mers within the top 100 TRBV2-1 clonotypes for each sample, with individual samples indicated by black boxes. Mean ± SD.

### Quantitative reduction in the response of TRBV2-1-expressing T cells in immunized IFNAR1^−/−^ animals

Next, we characterized the αβ T cell receptor (TCR) repertoire to assess whether T cell responses differ between IFNAR1^−/−^, IFNLR1^−/−^, and WT animals. Additionally, we aimed to determine if antibody responses to KLH are associated with clonal T-cell responses, potentially indicating T-dependent antibody responses. Both α and β chains were sequenced from T cells derived from the spleen at day 5 post-booster immunization from all three groups. Unimmunized chickens from each genotype were included to establish a baseline for the expression of TCRs. Interestingly, the TCR repertoires across genotypes and immunization groups were remarkably similar for both α and β chains, as indicated by the pairing of V-J gene segments (Supplementary Figs. 6 and 7). This suggests that the global repertoires are consistent across different genotypes after immunization. Given the greater diversity of β chains compared to α chains, resulting from the recombination of V, D, and J gene segments, we focused on β chains for the subsequent analyses of CDR3 clonotypes. CDR3 spectratypes exhibited a Gaussian-like distribution in both control and immunized chickens across all genotypes, suggesting that KLH does not induce major clonal expansions in the splenic TCR repertoire 5 days after booster immunization (Fig. 5a). This was confirmed by the analysis of clonotype frequencies, which predominantly consisted of rare clonotypes with only a few moderately expanded clonotypes observed in both control and immunized chickens (Supplementary Fig. 8). Although overall Vβ gene usage was similar in control and immunized chickens, TRBV2-1 was significantly less prevalent in KLH-immunized IFNAR1^−/−^ chickens than in WT immunized animals (Fig. 5b). This suggests a reduced response of TRBV2-1-expressing T cells to KLH in IFNAR1-deficient chickens. To investigate whether specific amino acid motifs were enriched in KLH-immunized WT but not in IFNAR1^−/−^ chickens, we analyzed the counts of 3-mer sequence motifs in TRBV2-1 CDR3 clonotypes, normalized by the total number of TRBV2-1 CDR3 sequences in each sample. Overall, the most prevalent 3-mers in immunized WT chickens were also present in immunized IFNAR1^−/−^ chickens, indicating that TRBV2-1 clonotypes in IFNAR1^−/−^ chickens are qualitatively similar to those in WT chickens (Fig. 5c). The observed differences are therefore more likely attributed to quantitatively reduced activation rather than distinct clonotype compositions.

#### Virus species- and strain-specific IFN responses

To investigate the role of type I and type III IFNs in the defense mechanisms against viral infections, an *in ovo* challenge experiment was performed using 11-day-old embryos (Supplementary Fig. 9). This experiment utilized four distinct viral strains: a laboratory-adapted human influenza A virus (H1N1), two low pathogenic avian influenza A virus strains (H3N1 and H9N2), and infectious bronchitis virus (IBV)- Beaudette strain, an important avian gamma coronavirus. A significant (P ≤ 0.05) increase in WSN33 (H1N1) viral titers in the allantoic fluid was observed in IFNAR1^−/−^ and IFNLR1^−/−^ embryos compared to WT embryos, indicating a crucial role for both type I and type III IFN signaling in restricting WSN33 replication. Conversely, H9N2 viral titers were significantly (P ≤ 0.05) reduced in IFNLR1^−/−^ embryos compared to IFNAR1^−/−^ and WT embryos, indicating a predominant role for type I IFN signaling in limiting H9N2 replication. However, no significant differences in viral titers were observed for H3N1 and IBV among the genotypes, suggesting potential strain-specific or virus-specific mechanisms governing the antiviral IFN response (Fig. 6a).

**Fig. 6.**
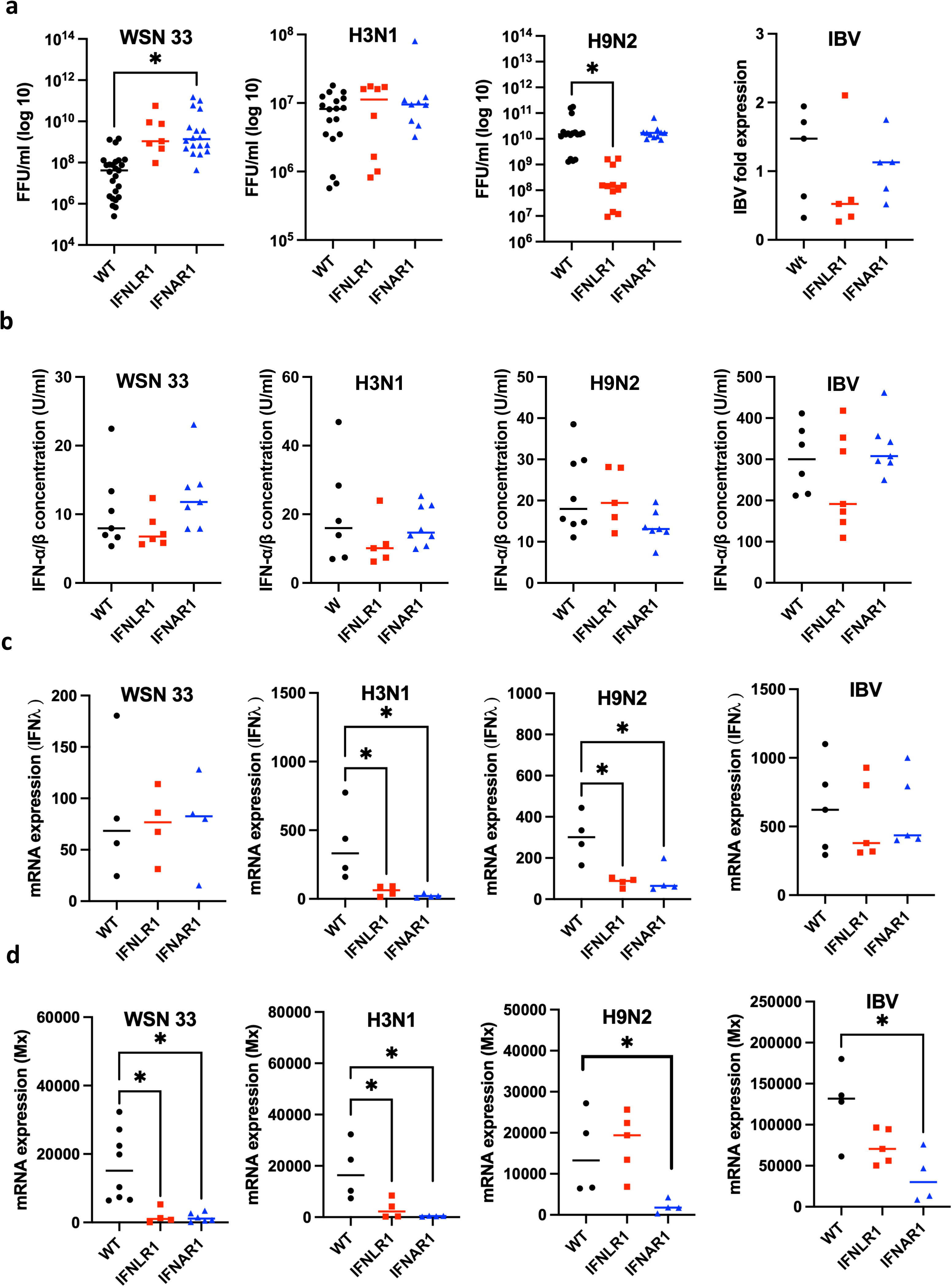
Viral titration, IFN-α/β concentration, IFN-λ, and Mx mRNA expression in WT, IFNLR1^−/−^ and IFNAR1^−/−^ embryos. **a**, embryonic day 11 chicken embryos were infected with 1000 FFU of influenza A virus strains (WSN33/H3N1/H9N2) and IBV- Beaudette strain, and allantoic fluid and chorioallantois membrane (CAM) were sampled 24 hours post-infection. The viral titers of WSN33, H3N1, and H9N2 were quantified by titration on MDCK cells. For the IBV, qPCR was used to measure the viral titer in the CAM, and the housekeeping gene 28S was used to normalize target gene expression. **b**, IFN-α/β concentration was quantified by titration of allantoic fluids on CEC-32 #511 cells. **c,** Relative mRNA expression of IFN-λ in the CAM. **d,** Relative mRNA expression of Mx in the CAM. For the qPCR study, RNA was isolated by Trizol and reverse transcripted with GoScript Transcription Mix, Random Primers, and qPCR was done via Go Taq. The WT control group was used as a calibrator for Mx and IFN-λ expression, and the housekeeping gene 18S was used to normalize target gene expression. Relative quantification of the target gene was performed using the 2^^ΔΔCT^ method. A no-template control was also included. Statistical differences between groups are indicated by asterisks (*), with significance at P ≤ 0.05.

**Fig. 7.**
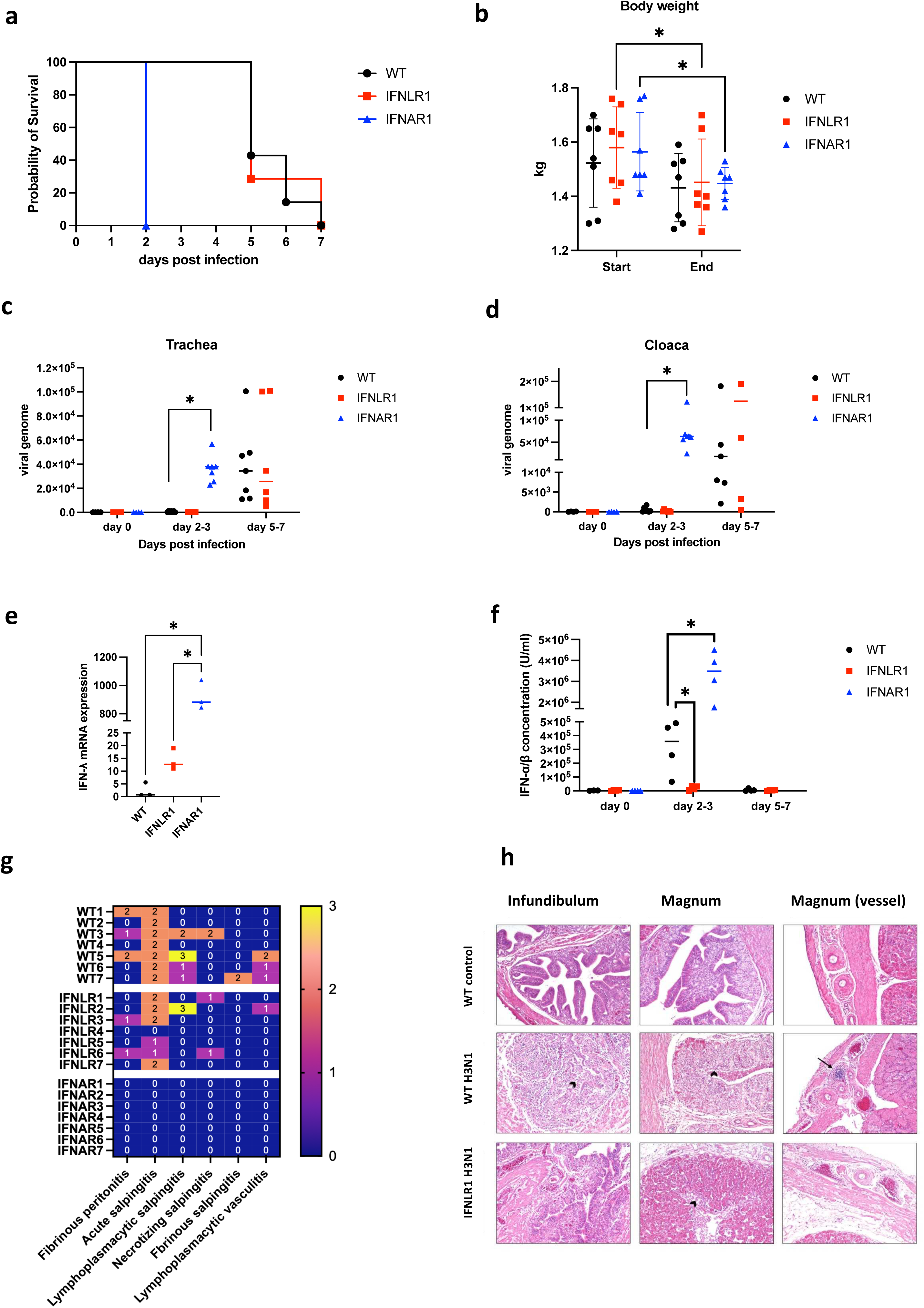
*In vivo* challenge of WT, IFNLR1^−/−^ and IFNAR1 ^−/−^ hens with a low pathogenic avian influenza A virus (H3N1). a, Probability of survival in the challenged hens. **b**, Chickens’ body weight (kg) was measured before the viral challenge and at the end of the experiment (date of animal euthanization). **c-d,** H3N1 titer in the tracheal and cloacal swabs using qPCR. The IFNAR1 hens were sampled on day 2 post-infection. WT and IFNLR1^−/−^ hens were sampled on day 3 and days 5 to 7 post-infection based on the days of euthanization. The housekeeping gene 28S was used to normalize target gene expression. **e,** Relative mRNA expression of IFN-λ in the spleen. Comparisons were designed to represent equivalent phases of disease progression. For panel e, IFNAR1^−/−^ hens at day 2 (high viral titer) were compared to WT and IFNLR1^−/−^ hens at day 5 (comparable high titer). **f,** Plasma IFN-α/β concentration was quantified by titration on CEC-32 #511 cells. For panel f, IFNAR1^−/−^ hens (day 2, high titer) were compared to WT and IFNLR1^−/−^ hens at day 3 (low titer) and days 5–7 (high titer), reflecting distinct infection phases. **g,** Scoring of lesions in the reproductive tract in all *in vivo* H3N1 challenged groups. The scoring is on a scale of 0-3, where 0 indicates normal and 3 maximum severities regarding lymphoplasmacellular salpingitis with mucosa atrophy. The IFNAR1^−/−^ hens were euthanized and scored on day 2 post-infection. WT and IFNLR1^−/−^ hens were euthanized and scored on days 5 to 7 post-infection based on the days of euthanization. **h,** Histological assessment of the reproductive tract on day 5 post infection. WT control: normal histological appearance of the infundibulum and magnum; WT H3N1: moderate acute inflammation of the infundibulum and magnum with hyperemia, edema, loss of epithelial cells (arrowheads), and perivascular infiltration of lymphocytes (arrow); IFNLR1^−/−^: milder acute inflammation of the infundibulum and magnum with hyperemia, edema, and loss of epithelial cells in the magnum (arrowhead). Hematoxylin-eosin (HE) stain; 100x, respectively. Statistical differences between groups are indicated by asterisks (**\*,** P ≤ 0.05).

#### Knockout of IFNAR1 and IFNLR1 did not affect IFN-α/β secretion *in ovo*

The initial virus-host interaction and IFN secretion occur independently of IFNAR1 and IFNLR1 receptors. However, subsequent IFN production is regulated through a feedback mechanism mediated by these receptors. Indeed, no difference in the concentration of IFN-α/β in the allantoic fluid of WT, IFNAR1^−/−^ and IFNLR1^−/−^ embryos was observed in response to viral infections *in ovo* (Fig. 6b). IFNλ mRNA expression in the chorioallantoic membrane (CAM) showed a significant (P ≤ 0.05) reduction in IFNAR1^−/−^ and IFNLR1^−/−^ compared to WT embryos challenged with low pathogenic avian influenza strains (Fig. 6c). Interestingly, the IBV- Beaudette strain infected embryos showed at least 10-fold higher induction of IFN-α/β secretion compared to the influenza strains used in this experiment (Fig. 6b).

### A predominant role of type I IFN in inducing the expression of an interferon-stimulated gene (Mx)

Mx expression in the CAM was studied to elucidate the contributions of type I and type III IFNs in inducing ISGs during viral infections. The CAM of chicken embryos was used as a model due to its high vascularization and functional similarity to avian respiratory and immunological tissues. CAMs isolated from IFNAR1^−/−^ embryos 24 hours post-infection exhibited a significant decrease (P ≤ 0.05) in Mx expression compared to WT embryos in all the tested viruses. In IFNLR1^−/−^ embryos, Mx levels in CAMs challenged with H9N2 and IBV-Beaudette strain remained comparable to Mx expression levels observed in WT embryos. These findings suggest a predominant role for type I IFN signaling in inducing the expression of ISGs, such as Mx, in response to H9N2 and IBV-Beaudette strain infections (Fig. 6d).

#### An excessive and unbalanced type I IFN response can harm host fitness

Survival probability experiments revealed that an overly strong IFN-α/β response may negatively impact host fitness more than the viral load. Consistent with this observation, measured IFN-α/β concentrations (Fig. 6b) were elevated in these embryos, supporting the link between high type I IFN responses and reduced survival. This was evident in IFNLR1^−/−^ embryos challenged with H9N2, which exhibited poor survival rates compared to WT and IFNAR1^−/−^ embryos despite having low viral titers. Similarly, the heightened type I IFN response likely explains the rapid death of IBV-challenged embryos within 24-36 hours post-infection. In contrast, embryos infected with influenza strains exhibited approximately tenfold lower IFN-α/β concentrations compared to IBV-challenged embryos and survived longer, lasting 4–5 days after infection (Supplementary Fig. 10). These results underscore the critical yet potentially dual role of the type I IFN response in controlling viral infection, as it may also contribute to immunopathology.

### Early appearance of symptoms and poor survival of IFNAR1^−/−^ hens challenged with H3N1

To gain deeper insight into the role of IFN-α/β and IFN-λ in the defense mechanism against low pathogenic avian influenza A virus strains (H3N1), an *in vivo* infection experiment was carried out (Supplementary Fig. 11). IFNAR1^−/−^ hens exhibited a rapid onset of severe symptoms at 48 hours post H3N1 infection, including conjunctivitis, diarrhea, mucus discharge, inflamed, red eyes, abnormal posture, and severely restricted movement compared to WT and IFNLR1^−/−^ hens, which only showed similar symptoms from day five onwards. All IFNAR1^−/−^ hens (n = 7) were euthanized by day 2 post-infection after reaching the humane endpoint to minimize suffering. In comparison, four WT hens were euthanized on day 5, two on day 6, and one on day 7 post-infection. Similarly, five IFNLR1^−/−^ hens were euthanized on day 6 and two on day 7 post-infection (Fig. 7a). Because humane endpoints were reached at different days for each genotype, body weight changes were analyzed within genotypes, and data are presented as mean ± SD (Fig. 7b). There was a significant reduction (P ≤ 0.05) in the body weight of IFNAR1^−/−^ and IFNLR1^−/−^ hens from the start to the end of the experiment. In contrast, while WT hens also showed a reduction in body weight by the end of the experiment, the change was not statistically significant (Fig. 7b).

### Efficient H3N1 replication in IFNAR1^−/−^ hens

In IFNAR1^−/−^ hens, the H3N1 virus titer increased significantly (P ≤ 0.05) by day 2 post-infection in both the trachea and cloaca, indicating the limited ability of the immune system to restrict viral replications in these birds. In contrast, WT and IFNLR1^−/−^ hens maintained low H3N1 titers until day 5 post-infection, with levels rising similar to those in IFNAR1^−/−^ hens (Fig. 7c,d).

### Increased plasma concentration of IFN-α/β and spleen expression of IFN-λ in IFNAR1^−/−^ hens

In IFNAR1^−/−^ hens, we observed an increase (P ≤ 0.05) in the mRNA expression of IFN-λ, suggesting a compensatory mechanism in the absence of type I IFN signaling in the spleen where IFN-λ receptors are expressed (Fig. 7e). For Fig. 7e, comparisons were designed to represent equivalent phases of disease progression: IFNAR1^−/−^ spleens at day 2 post infection (high viral titer) were compared with

WT and IFNLR1^−/−^ spleens at day 5 post infection, when viral titers were comparably high. In IFNAR1^−/−^ hens, the concentration of IFN-α/β showed around 10 times higher (P ≤ 0.05) levels than in WT and IFNLR1^−/−^ hens at day 2-3 post-infection, indicating uncontrolled secretion of IFN-α/β in IFNAR1^−/−^ hens (Fig. 7f). For Fig. 7f, IFNAR1^−/−^ hens at day 2 (high titer) were compared with WT and IFNLR1^−/−^ hens at day 3 (low titer) and days 5–7 (high titer), thereby sampling corresponding disease stages across genotypes.

#### IFN-λ signaling promotes oviduct inflammation in H3N1-infected chickens

The H3N1 avian influenza A virus demonstrates a distinct tropism for the hen’s oviduct, causing salpingitis and peritonitis associated with a severe decline in egg production. While young birds showed minimal clinical signs, adult layers exhibited higher mortality and decreased egg production in the second week post-infection^29^. To evaluate the impact of type I and type III IFN on the pathogenesis of H3N1, we evaluated oviduct inflammation, the site where H3N1 establishes the infection. The oviduct inflammation was scored on a magnum grade scale of 0-3 (0 indicating an intact oviduct and 3 indicating severe inflammation)^30^. The results revealed a magnum grade 0 for the oviduct collected from IFNAR1^−/−^ at day 2 post-infection. The IFNLR1^−/−^ hens predominantly exhibited magnum grade 1, indicating mild lymphoplasmacellular salpingitis with normal mucosal height. In contrast, WT hens mostly displayed magnum grade 2, suggesting moderate lymphoplasmacellular salpingitis with prominent atrophy of the mucosa (Fig. 7g). Histological analysis of the oviduct on day 5 post-infection revealed moderate acute inflammation in the infundibulum and magnum with hyperemia, edema, epithelial cell loss, and perivascular infiltration of lymphocytes in WT hens challenged with H3N1 compared to WT negative controls. In contrast, IFNLR1^−/−^ hens displayed milder inflammation in these parts of the oviduct (Fig. 7h). These results indicate a pro-inflammatory role of IFN-λ in H3N1 infection in the oviduct.

### Increased expression of pro-inflammatory cytokines in the spleen of IFNAR1^−/−^ hens

In spleens sampled two days after infection, IFNAR1^−/−^ hens showed a significant (P ≤ 0.05) increase in the mRNA expression of several pro-inflammatory cytokines, including IL-1β, IL-2, IL-6, IL-8, IL-12, and IL-22. This increase was observed compared to both WT and IFNLR1^−/−^ hens, whose spleens were sampled five days post-infection. Notably, IL-17A expression did not show a significant difference between the groups (Supplementary Fig. 12a). The elevated levels of IL-2 and IL-12 point to the activation of the Th1 pathway^31^. The upregulation of IL-1 and IL-6 suggests an induction of the Th22 pathway, known to stimulate Th22 cells, a recently characterized CD4^+^ T cell lineage, to release IL-22 but not IL-17A^32, 33^. This would lead to inflammation and tissue damage as overexpression of IL-22 is associated with inflammatory tissue pathology^34^. Also, there was no change in the expression of genes associated with the anti-inflammatory pathway and regulatory T cells (FOXP3, IL-10, TGF-β), indicating a lack of a compensatory anti-inflammatory mechanism to control the inflammation in IFNAR1^−/−^ hens. Additionally, there was a significant (P ≤ 0.05) decrease in the expression of IL-4 and IL-5 in IFNAR1^−/−^ hens, which indicates lower Th2 pathway activity in these hens (Supplementary Fig. 12a).

### Impairment of negative feedback mechanisms regulating IFN-α/β secretion and low antiviral state in the spleen of IFNAR1^−/−^ hens

A gene expression study was performed on the spleen to gain deeper insights into the alterations in H3N1-host cell interactions and modulation of IFN signaling in IFNAR1^−/−^ hens. mRNA expression analysis of genes related to H3N1-host cell interaction, pro-inflammatory cytokines, and IFN induction (TLR3, MDA5, MyD88, NFkB, and IRF7) showed no significant differences between the groups (data not shown), suggesting that these pathways function normally in both WT and IFNAR1^−/−^ cells (Fig. 8 a,b). This could also be attributed to the fact that the spleen used in this analysis was collected from WT and IFNLR1^−/−^ at day 5 post-infection, a time when the viral load was high in these animals, and this pathway was active, similar to the IFNAR1^−/−^ hens. Viral sensation by the host cell and activation of IFN-α/β secretions occur in various cell types expressing TLRs, including immune cells, epithelial cells, fibroblasts, and endothelial cells^35, 36^. The interaction with the IFNAR1 receptor is needed for the secreted IFN-α/β to mediate its function. The knockout of IFNAR1 led to a significant decrease in this group’s antiviral state. Relative mRNA expression of antiviral genes induced by IFN, including viperin, ISG12, IFITM5, PKR, and STAT1, reflects significantly (P ≤ 0.05) lower antiviral state in the IFNAR1^−/−^ group as compared to WT and IFNLR1^−/−^ hens. These results indicate that IFN-α/β can compensate IFN-λ to induce an adequate antiviral state during H3N1 infection. However, IFN-λ can not compensate for IFN-α/β in the spleen (Supplementary Fig. 12b). In WT cells, negative feedback mechanisms mediated by suppressor of cytokine signaling 1 (SOCS1) and Src Homology 2-containing Phosphatase 2 (SHP2) regulate the secretion of IFN and associated inflammation once an adequate antiviral state is achieved. In IFNAR1^−/−^ cells, a low antiviral state, and high viral titers likely prevent the efficient activation of the negative feedback loop, as indicated by the reduced expression of SOCS1 and SHP2 (Supplementary Fig. 12c). Consequently, this leads to the continuous and significant secretion of IFN-α/β (Fig. 8).

**Fig. 8.**
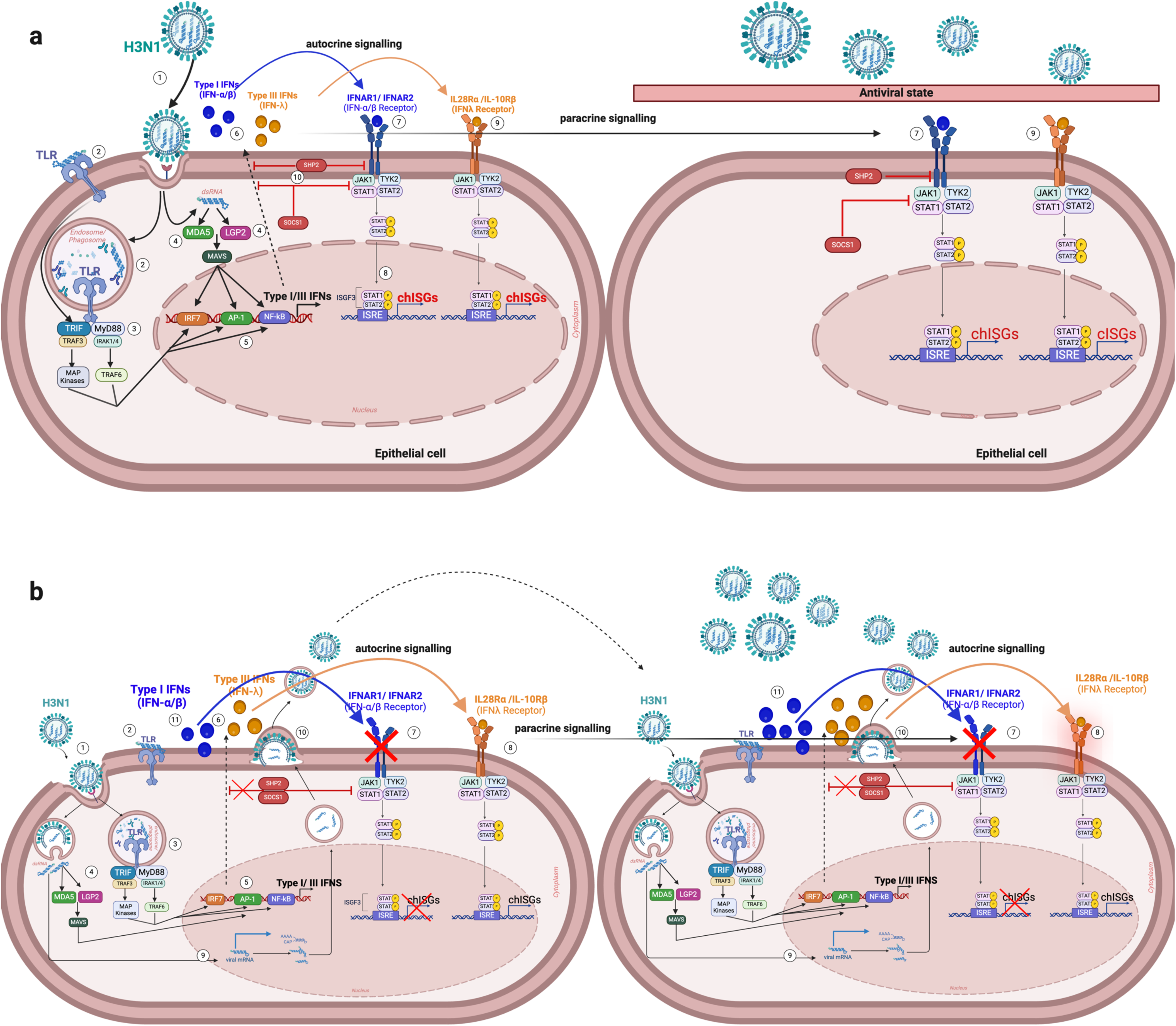
Schematic representation of H3N1 host cell interaction and IFN signaling pathway in WT and IFNAR1^−/−^ at day 2 post-H3N1 challenge. a, Immune signaling pathway in WT epithelial cells during an H3N1 viral infection. The cleaved hemagglutinin (HA) proteins of H3N1 bind to sialic acid receptors on the surface of epithelial cells, facilitating viral entry through endocytosis (1). The viral RNA inside the endosome is recognized by toll-like receptors (TLRs) (2), leading to the activation of the MYD88 signaling pathway (3) and melanoma differentiation-associated protein 5 (MDA5) and/or laboratory of genetics and physiology 2 (LGP2) (4). In the MDA5/LGP2 pathway, mitochondrial antiviral-signaling proteins (MAVS) are activated. The activation of MYD88 leads to the recruitment of IRAK1/4 and TRAF6/3, which then stimulate the transcription factors nuclear factor kappa-light-chain-enhancer of activated B cells (NF-κB), interferon regulatory factor 7 (IRF7) and activating protein 1 (AP-1) (5). These transcription factors induce the production of pro-inflammatory cytokines and type I and type III IFNs (6). Type I IFN (IFN-α/β) binds to its receptors (IFNAR1/2) by autocrine and paracrine signaling (7). This binding activates the JAK-STAT pathway, leading to the phosphorylation of signal transducers and activators of transcription (STATs). These STATs translocate to the nucleus and activate the transcription of interferon-stimulated response elements (ISREs). The activation of ISREs drives the expression of interferon-stimulated genes (ISGs), establishing a robust antiviral state that effectively halts viral replication and spread within the cell (8). Similarly, type III IFN (IFN-λ) contributes to the antiviral state by binding to its receptors (IL-28Rα /IL-10Rβ) and activating the JAK-STAT pathway in an autocrine or paracrine manner (9). The immune response is tightly regulated by negative feedback mechanisms, including suppressor of cytokine signaling (SOCS1) and Src Homology 2-containing Phosphatase 2 (SHP2), which inhibit further production of IFNs and prevent excessive inflammatory responses (10). This response ensures that any initial viral replication is quickly suppressed, protecting the cell from significant infection and excessing immune responses that could lead to cytokine storm. The viral replication is significantly halted since the cells are in an antiviral state. **b,** Immune signaling pathway in IFNAR1^−/−^ epithelial cells. Virus entry to the cell, recognition of viral RNA by the host cell, and induction of pro-inflammatory cytokines and IFNs happen in a mechanism similar to the WT cell (steps 1-6). However, without IFNAR1, IFN-α/β cannot bind to its receptors, disrupting downstream signaling and feedback inhibition, leading to unregulated IFN production and poor antiviral state (7). Higher IFN-λ is produced to compensate for the low antiviral state but cannot compensate for the absence of IFN-α/β (8). This makes the infected and neighboring cells easy targets for the H3N1 virus that can replicate quickly without resistance. Once inside the cell in the endosome, the viral RNA is released and transported into the nucleus for replication and transcription (9). Newly formed viral particles assemble, bud off, and are released to infect additional cells (10). In the absence of IFNAR1 and an antiviral state, the autocrine feedback loop mediated by SOCS1 and SHP2 is disrupted, and thus, the production of IFN-α/β is continuously increasing (11). The extensive pro-inflammatory cytokines and IFN production exacerbate inflammatory responses and cell death.

## Discussion

Interferons are crucial components of the innate immune system, playing a pivotal role in host defense against viral, bacterial, and other pathogenic insults. In mammals, the potent antiviral activities induced by IFNs have rendered them attractive therapeutic agents for bolstering the immune response against infections and diseases^37, 38, 39^. Notably, both IFN-α/β and IFN-λ have demonstrated potential as adjuvants to enhance the efficacy of mucosal vaccines, particularly against respiratory viruses like influenza in murine models^40, 41^. Developing novel therapeutic approaches to combat and eradicate viral and bacterial infections in poultry necessitates a comprehensive understanding of the mechanisms of action of type I and type III IFNs. To investigate this further, we have generated type I and type III IFN receptors knockout chickens to elucidate their potential roles in controlling avian influenza and other avian viral diseases. The IFNAR1^−/−^ and IFNLR1^−/−^ chickens exhibited normal growth and development compared to their WT counterparts. Flow cytometry analysis of the immune cell population in blood, spleen, and plasma concentrations of immunoglobulins indicate that IFN-α/β plays a crucial role in regulating innate immune cells, T cell subsets, and their contribution to antibody production under homeostatic conditions.

Both IFNAR1^−/−^ and IFNLR1^−/−^ chickens exhibited impaired antibody production in response to KLH immunization. Phagocytosis and antigen processing are augmented in a properly functioning interferon system, contributing to efficient antibody production in chickens^42^. In avian species, macrophages and B cells are well-characterized antigen-presenting cells (APCs) that express MHCII molecules and present antigens to CD4^+^ T helper cells, leading to antibody production^42, 43^. On day 5 after booster immunization, the spleens of IFNAR1^−/−^ chickens exhibited fewer B cells, macrophages, and MHCII^+^ cells, including MHCII^+^ B cells and MHCII^+^ macrophages, than other genotypes. Loss of IFN sensing led to reduced regulatory T cell selection and reduced TCR repertoire diversity in a murine model^44^.

*In ovo* challenge of both IFNAR1^−/−^ and IFNLR1^−/−^ embryos revealed a complex interplay between chicken IFN signaling and viral pathogenesis, demonstrating that susceptibility to infection and subsequent immune responses highly depend on viral species and strain. Increased susceptibility and poor restrictions of viral replication in IFN receptor knockout embryos were observed only for WSN33 (H1N1) strain. At the same time, no significant differences in viral titer were noted between the groups in response to H3N1. The case of low pathogenic avian H9N2 virus is particularly intriguing. Despite causing comparatively few deaths, H9N2 has garnered significant attention due to its potential genetic exchange with emerging zoonotic influenza viruses such as H7N9 and H10N8 subtypes^45, 46^. Interestingly, we observed that reduced H9N2 titers in IFNLR1^−/−^ did not correlate with improved embryo survival compared to other genotypes. This finding highlights the critical role of chicken IFN-λ as anti-inflammatory cytokines in maintaining immune balance during the antiviral defense against H9N2.

Because viral titers and type I IFN concentrations were measured in the allantoic fluid, while Mx expression reflected responses in the chorioallantoic membrane (CAM), these analyses delineate distinct layers of interferon regulation. The CAM’s mixed cellular composition allows systemic-type I IFN signaling through vascular and mesodermal layers, whereas type III IFN activity via IFNLR1 is concentrated at the epithelial surface^11, 47^. In our assays, Mx expression was measured at 24 hours post-infection, a time when type I IFN response may be highly active and capable of compensating for loss of type III IFN even in response to viruses with a predominant or prolonged epithelial phase like H9N2 and IBV^48, 49^. This redundancy may explain why type I IFN signaling alone was sufficient to drive robust Mx induction in IFNLR1 knockout embryos for these viruses, while type III IFN remains essential for maximal Mx induction in infections that spread to both compartments (e.g., WSN33, H3N1)^29, 50^. Thus, the observed differential impacts of receptor knockout reflect both spatial compartmentalization and the timing of IFN responses, rather than a lack of type III IFN function in epithelial antiviral defense

Although H3N1 is classified as a low pathogenic avian influenza A virus due to its monobasic hemagglutinin cleavage site, it has shown high virulence in adult layers, causing severe clinical infections and mortality^29^. Researchers revealed that H3N1 can rapidly replicate and disseminate systemically in chickens due to a neuraminidase mutation that activates plasminogen-mediated hemagglutinin^51^. This mechanism mimics highly pathogenic strains and raises concerns about potential zoonotic transmission, particularly given the history of H3 subtypes in human infections. The *in vivo* H3N1 infection in WT, IFNLR1^−/−^ and IFNAR1^−/−^ hens revealed significant differences in disease progression and immune responses, emphasizing the critical role of type I IFN signaling in the defense mechanism against this H3N1 strain. In contrast, in mammalian systems, disruption of both type I and type III IFN pathways is generally required to produce comparable effects^18, 19^. These findings suggest that type I IFNs may play a more dominant and less redundant role in the response to the H3N1 strain and point to species-specific differences in interferon biology that warrant cautious comparison with mammalian systems.

Previous studies suggested that H3N1 replication in the oviduct is a critical factor in the onset of clinical disease and increased viral excretion^29^. Our current study provides new insights into H3N1 pathogenesis. The IFNAR1^−/−^ group displayed severe clinical symptoms and higher viral titers despite having intact oviducts and egg production. This observation suggests that H3N1 can establish infection and replicate independently of oviduct involvement, indicating a more complex pathogenic mechanism than previously thought. IFN-λ, initially described as an anti-inflammatory counterpart to type I IFN, is highly expressed by epithelial cells, which line the avian oviduct^52^. Recent studies have revealed that mammalian IFN-λs can exert complex effects on innate and adaptive immunity, sometimes promoting inflammation in specific contexts^20, 21, 53^, which agrees with the lower and delayed severity of oviduct infection and tissue damage in IFNLR1^−/−^ compared to the WT hens. This observation suggests a potential pro-inflammatory role for IFN-λ in the context of H3N1 infection in the avian oviduct. Similarly, mammalian type III IFNs can disrupt the lung epithelial barrier upon viral recognition, showing detrimental effects during chronic viral infection^20^. Moreover, IFN signaling, particularly IFN-λ, impairs lung repair by inducing p53 and inhibiting epithelial proliferation and differentiation in a mouse model of influenza infection^21^.

SOCS1 and SHP2 are pivotal regulators of IFN signaling, playing a crucial role in maintaining a delicate balance between effective immune responses and homeostasis^36, 54^. The lack of negative feedback mediated by SOCS1 and SHP2 in the IFNAR1^−/−^ hens, due to the absence of an antiviral state and increased viral spread, led to a continuous and significant secretion of IFN-α/β. These results agree with several studies that reported IFNAR1^−/−^ mice produce high levels of circulating IFNs. This phenomenon has been attributed to the lack of an IFNAR1 receptor to bind and internalize type I IFNs, reduced clearance of IFNs from circulation, and continued production of IFNs by cells in response to ongoing stimuli, resulting in their accumulation^24, 55^.

Type I IFNs are crucial in moderating leukocyte infiltration and the inflammatory cascade. However, disrupting the IFNAR1 receptor can lead to a dysregulated cytokine response and excessive inflammation^56, 57^. In IFNAR1^−/−^ hens, eliminating type I IFN signaling and unaffected expression of inflammatory cytokines allows the H3N1 virus to spread further. The increased presence of viral components in the system continuously amplifies inflammatory cytokine production, potentially leading to a cytokine storm. Our observations support this hypothesis, as we found increased mRNA expression of pro-inflammatory cytokines in the spleen of IFNAR1^−/−^ hens, which indicates a clear activation of Th1 and Th22 pathways in these birds^31, 32, 33^. The simultaneous activation of multiple T helper cell subsets likely contributed to the rapid onset of severe symptoms observed in IFNAR1^−/−^ hens. Over-induction of Th1 and Th22 responses can lead to significant tissue damage and chronic inflammation, potentially increasing the risk of autoimmune-like reactions when left unregulated. Regulatory T cells (Tregs) normally modulate these responses by producing anti-inflammatory cytokines such as IL-10 and TGF-β^58^. However, our findings of unchanged expression of genes associated with the Tregs (FOXP3, IL-10, TGF-β) suggest that Treg activation was insufficient in the IFNAR1^−/−^ group.

Previous research on murine models has demonstrated that reduced inflammation can confer resistance to influenza infection, even in elevated viral loads^59, 60^. These observations suggest that the inflammatory response triggered by the virus may play a more crucial role in influenza pathology than the viral burden itself. Based on this understanding, we propose that the heightened inflammatory response observed in IFNAR1^−/−^ hens is a key factor in the increased pathology associated with H3N1 infection. Our findings indicate that the IFNAR1 receptor is critical for preserving the delicate balance between potent antiviral immunity and the potentially harmful consequences of unrestrained inflammatory responses.

The present study uncovered the critical roles of type I and III IFNs on antibody production and baseline immune cell populations in chickens. Furthermore, these knockout models revealed that type I and type III IFNs play distinct and crucial roles in modulating viral pathogenesis, immune responses, and tissue-specific effects during avian influenza infection. The roles of these IFNs are virus strain-specific, with type I IFN showing particular importance in early defense against H3N1 avian influenza. By dissecting the complexities of avian interferon responses, these chicken models are critical for developing more effective strategies to control avian viral infections, including avian influenza, impacting veterinary medicine and zoonotic disease control.

## Material and Methods

### Animals

Specific pathogen-free (SPF) hatching eggs from white Lohman selected leghorn (LSL) were obtained from ValoBioMedia GmbH (Osterholz-Scharmbeck, Germany). These chickens were housed at the TUM Animal Research Center (TUM School of Life Sciences, Weihenstephan). They were provided with water and commercial feed *ad libitum*. All animal experiments were conducted following current laws and approved by the government of Upper Bavaria (experiment licenses: ROB-55.2-2532.Vet_02-20-13 & ROB-55.2-2532.Vet_02-23-100).

### Generation of IFNAR1 and IFNLR1 KO primordial germ cells (PGCs)

LSL PGCs were derived from the blood of embryonic vasculature at stages 13-15, as described earlier^61^. These PGCs, derived from male embryos, were cultured at 37°C in a 5% CO2 environment using modified KO-DMEM^62^. Guide RNAs targeting the coding region within exon 1 of the IFNAR1 chain of the IFN-α/β receptor and the epithelium-specific chain (IFNLR1) of the IFN-λ receptor (Supplementary Table 1) were designed and cloned separately into the PX458-Cas 9 vector (Addgene, USA), which contains eGFP as a selectable marker. A single-stranded oligodeoxynucleotide (ssODN) (Supplementary Table 1) was synthesized by Integrated DNA Technologies (IDT, Zellik, Belgium) to serve as a repair template for the IFNAR1 KO. The PGCs were transfected with the targeting vector (PX458-sgRNA) and sorted based on eGFP fluorescence with fluorescence-activated cell sorting (FACS) as previously described^63^. Selected eGFP-positive PGCs were plated on a 48-well plate to expand single-cell clones. Sanger sequencing was employed to identify the desired genetic modification.

### Generation of IFNAR1^−/−^ and IFNLR1^−/−^ chickens

After successful clonal selection, PGCs were genotyped to identify clones with a knockout. These clones were used to generate chimeric roosters. For IFNAR1, the INDEL was pre-designed using CRISPR-mediated HDR technology, resulting in a 7-bp deletion that produced a stop codon after 21 bp. For IFNLR1, the clone with the maximum deletion (28 bp) was selected as it allowed for the easy design of an IFNLR1 KO-specific PCR genotyping assay and produced a stop codon after 87 bp (Fig.1 a,b). The knockout PGC clones were injected into the vasculature of 65-hour-old embryos and incubated until the hatch. Upon sexual maturity, sperm was collected, and germline transmission was evaluated as previously described^64, 65^. Germline-positive chimeric roosters were bred with WT hens to obtain heterozygous animals. IFNAR1 and IFNLR1 homozygous birds were generated by crossing the heterozygous birds (IFNAR1^+/−^) and (IFNLR1^+/−^), respectively. Following the generation of different transgenic lines, the birds were monitored until sexual maturity for any harmful phenotypes, such as impacts on weight gain and reproductive capabilities. Additionally, homozygous roosters and chickens were bred within each line to test for growth rate and the ability to produce fertile sperm or eggs.

### Genotyping of genetically modified chickens

For the genotyping of IFNAR1 KO, TaqMan assay was designed and performed using 5x HOT Firepol Probe qPCR Mix Plus (Solis Biodyne, Estonia). Detailed information about the sequence of the 1263-IFNAR1-For and 1264-IFNAR1-Rev primers (IDT, Zellik, Belgium) as well as 1291- IFNAR1 MUT-probe_HEX and 1262- IFNAR1 WT-probe_FAM probes (Eurofins, Ebersberg, Germany) can be found in Supplementary Table 2. The following cycler conditions were used: Pre-read stage at 60°C for 30 sec, hold stage at 95°C for 15 min, followed by 40 cycles of 95 °C for 15 sec, and 60 °C for 1 min, and a post-read stage of 60 °C for 30 sec. The genotyping of IFNLR1 KO was done via endpoint PCR using Firepol DNA Polymerase (Solis Biodyne, Estonia) employing the IFNLR1 primers shown in Supplementary Table 2. The following PCR conditions were used: initial denaturation at 95°C for 3 min, followed by 40 cycles of 95 °C for 15 sec, 59 °C for 30 sec, 72 °C for 15 sec, and a final extension step of 72 °C for 5 min. The WT sample produced an amplicon size of 250 bp, the homozygote produced an amplicon size of 222 bp, and the heterozygote sample produced amplicon sizes of both 250 bp and 222 bp.

### Western blot and RT-PCR to examine successful knockout of IFNAR1/IFNLR1

To confirm the deletion of IFNAR1, a western blot experiment was conducted to study the expression of Mx protein in chicken CEFs treated with recombinant IFN-α. CEFs were isolated following a previously established protocol from day 10-old embryos, with genotype confirmation through blood collection from a 0.5 cm² window in the eggshell that allowed access to the embryonic vasculature^66^. The CEFs were cultured and maintained as described earlier^63^. WT, IFNAR1^+/−^, and IFNAR1^−/−^ CEFs (n = 3) were cultured in a 6-well plate and stimulated with recombinant IFN-α (500 U/ml) for 12 hours. A control group of CEFs was treated with CEF medium without recombinant IFN-α. Following the stimulation period, both control and treated CEF cells were trypsinized and collected for SDS-PAGE and Western Blot analysis to study the expression of chicken Mx protein (75 kDa) and the control protein chicken β-actin (42 kDa). Mouse anti-human monoclonal IgGκ antibodies targeting MxA (M143; Merck, Germany) and mouse anti-chicken monoclonal IgG2b antibodies targeting β-actin (Thermo Fisher Scientific, Germany) were used as primary antibodies as described earlier^67^. HRP-conjugated donkey anti-mouse IgG was used as the secondary antibody (Thermo Fisher Scientific, Germany). Blots were developed with homemade ECL substrate, and the signal was detected using the Vilber Lourmat Fusion Fx system (Eberhardzell, Germany). Additionally, an experiment examined the absence of viral resistance in IFNAR1^−/−^ embryos compared to WT embryos after stimulation with recombinant IFN-α. Day 10-old chicken embryos were stimulated with 1.5 × 10^5^ U of recombinant IFN-α in 100 µl PBS 12 hours before and at the time of infection, while control groups did not receive IFN-α stimulation. All groups of embryos (four embryos per group) were infected with 1000 FFU of WSN33, and the viral load in the allantoic fluid was analyzed 24 hours post-infection by titration on the Madine-Darby canine kidney (MDCK) cells, as described earlier^67^.

To confirm the deletion of IL-28Rα (IFNLR1), various tissues (spleen, liver, lung, intestine, bursa, heart, and cecum) were collected from 18-day embryos of WT, IFNLR1^+/−^, and IFNLR1^−/−^. To ensure that the expression of IL-28Rα is induced, these embryos were challenged with 1000 FFU of H3N1 strain 12 hours before organ collection. The organ samples were rinsed with PBS and homogenized using a SpeedMill Homogenizer (Analytik Jena). After genotyping the embryos, RNA was isolated using the Trizol method (VWR, Germany)^68^, followed by cDNA synthesis with the GoScript Reverse Transcription Mix (Promega, USA). PCR was conducted using β-actin primers (277-FW & 278-RV) and IL-28Rα primers (1642-FW & 1643-RV) (Supplementary Table 3). The PCR was performed with Firepol DNA Polymerase (Solis Biodyne, Estonia) according to the manufacturer’s instructions. The PCR conditions were as follows: initial denaturation at 95°C for 3 min, followed by 40 cycles for β-actin or 35 cycles for IL-28Rα of 95°C for 30 seconds, 56°C for β-actin or 62°C for IL-28Rα for 30 seconds, and 72°C for 25 seconds. A final extension step was carried out at 72°C for 5 min. Nuclease-free water (NFW) was used as a negative control for both. A 1.5% Tris-borate-EDTA (TBE) gel was used to visualize the β-actin amplicon at 300 bp, and a 2% TBE gel was used to visualize the 108 bp amplicon for IL-28Rα.

### Experimental design

Various *in vitro*, *in ovo*, and *in vivo* experiments were conducted to characterize the IFNAR1^−/−^ and IFNLR1^−/−^ genetically modified lines. The first experiment aimed to study the impact of type I and III IFNs on growth development, immune response, and cecal microbiota. Heterozygous birds of each line were bred separately, resulting in WT, heterozygous, and homozygous birds for each line. At least 12 animals per group of WT, IFNAR1^−/−^, and IFNLR1^−/−^ were included in the study. Six chicks per group were euthanized at the age of 1 month to examine differences in immune cell populations in blood and spleen. Cecal contents from IFNLR1^−/−^ chickens and their WT siblings were collected for microbiota analysis. In the second experiment, the ability of IFNAR1^−/−^ and IFNLR1^−/−^ chickens to respond to immunization and secrete antibodies was investigated. Keyhole limpet hemocyanin (KLH) was used as a model T cell–dependent antigen to assess adaptive immune and antibody responses, providing a robust system for evaluating B cell activation and humoral immunity in knockout models^69^. Five-week-old WT, IFNLR1^−/−^, and IFNAR1^−/−^ chicks were immunized via intramuscular injection with 300 µg KLH, mixed 1:1 with incomplete Freund’s adjuvant. A booster injection of 300 µg KLH, mixed 1:1 with Freund’s incomplete adjuvant, was administered two weeks after the initial immunization. Plasma samples were collected from all groups before immunization (day 0), on day five post-primary immunization (PP), and on days 3 and 5 post-booster immunization (PB). The total concentration of IgM, IgY, and levels of KLH antigen-specific IgM and IgY were assessed using ELISA. Peripheral blood mononuclear cells (PBMCs) were isolated on day three post-PP, and splenocytes were isolated on day 5 PB for FACS analysis of immune and MHC-positive cells. Spleen samples were used to study TCR repertoire. At least six animals per group were included (Supplementary Fig. 3).

In the third experiment, an *in ovo* study was conducted to examine the effects of IFNAR1 and IFNLR1 knockout on viral replication, survival probability, Mx and IFNλ mRNA expression, as well as IFN-α/β concentrations in WT, IFNLR1^−/−^, and IFNAR1^−/−^ embryos (Supplementary Fig. 9). Fertilized eggs from a crossing of heterozygous roosters and hens of each line was collected and incubated until embryonic day 11 (ED11). The eggs were infected with 1,000 FFU of different influenza strains (WSN33(H1N1)/H3N1/H9N2) and infectious bronchitis virus (IBV)- Beaudette strain separately, as described previously^70^. The experiment was performed blindly, and the genotype of embryos was determined, and only WT and homozygous embryos were selected for further analysis. Details about the viral strains used in this study are provided in Supplementary Table 4. Twenty-four hours post-infection, the allantois fluid and breast tissue were collected for viral titration and genotyping, respectively. The CAM was also sampled 24 hours post-infection for gene expression studies. As previously described, the viral titers of WSN33, H3N1, and H9N2 were quantified by titration on MDCK cells^67^. For IBV, qPCR was used to measure the viral expression in the CAM (Supplementary Table 3). The relative mRNA expression of Mx and IFNλ was measured in the CAM. IFN-α/β concentration was quantified by titration of allantoic fluids on CEC-32 #511 reporter cells expressing firefly luciferase under the control of the chicken Mx promoter^71^. Plasmid #511 contains resistance to the antibiotic Geneticin (G418, 250 μg/ml) to keep the cells under selection pressure.

In the fourth experiment, the antiviral and immunomodulatory properties of type I and type III IFNs in the defense against H3N1 were characterized under *in vivo* conditions (Supplementary Fig. 11). The experimental groups consisted of WT, IFNLR1^−/−^, and IFNAR1^−/−^ hens, with 10 hens in each group. Seven hens per group were infected with H3N1, while the remaining three were negative controls. The hens, 27 weeks old and actively laying eggs, received an infectious dose 10^6^ FFU of H3N1 in 0.2 ml PBS per hen via nasal and tracheal routes on day 0. The H3N1 virus (A/Chicken/Belgium/460/2019) was provided by Dr. Joris Pieter De Gussem (Poulpharm BV). All IFNAR1^−/−^ hens (n = 7) were euthanized by day 2 post-infection due to early and severe symptoms. In comparison, four WT hens were euthanized on day 5, two on day 6, and one on day 7 post-infection. Similarly, five IFNLR1^−/−^ hens were euthanized on day 5 and two on day 7 post-infection based on symptom severity. Cloacal and tracheal swabs for viral RNA loads by RT-qPCR^72^ and blood samples for quantifying plasma IFN-α/β concentrations are taken on day 0 (all groups), day 2 (only IFNAR1^−/−^), day 3 (WT and IFNLR1^−/−^), and day 5-7 post-infection (WT and IFNLR1^−/−)^ based on the appearance and severity of symptoms. Blood plasma IFN-α/β concentrations were quantified using luciferase assay as described in experiment 3. Spleen samples were collected from all the infected animals for gene expression study. The oviducts were also collected from all the hens to study the impact of type I and type III IFN on the pathogenesis of H3N1 (Supplementary Fig. 11).

### Isolation of peripheral blood and splenic mononuclear cells

PBMCs and mononuclear spleen cells were isolated using Histopaque®-1077 density gradient centrifugation (Sigma-Aldrich, Darmstadt, Germany). Collected blood was diluted 1:1 with PBS and layered onto Histopaque®-1077, followed by centrifugation at 400 × g for 20 min at room temperature without brake. The mononuclear cell layer was collected, washed twice with PBS, and resuspended in FLUO-Buffer (1x PBS + 1%BSA + 0.01% NaN3). For spleen samples, tissues were aseptically dissociated through a 100 μm cell strainer into PBS to obtain single-cell suspensions. After centrifugation, cells were layered onto Histopaque®-1077 and processed as described for PBMCs to obtain the mononuclear fraction.

### Flow cytometry

Flow cytometric analyses were performed on PBMCs and mononuclear spleen cells isolated as described above. The detailed panel of primary and secondary antibodies employed for the identification and characterization of B cells, monocytes, γδ T cells (TCR1), αβ T cells (TCR2 + TCR3), and CD4-CD8 T cell subsets in experiment 1 is provided in Supplementary Table 5. Similarly, the antibody panel utilized for the analysis of B cells, monocytes, γδ T cells (TCR1), αβ T cells (TCR2 + TCR3), major histocompatibility complex class I (MHCI), and class II (MHCII), MHCII+ B cells, and MHCII+ monocytes in experiment 2 are detailed in Supplementary Table 5. Briefly, 5 x 10^6^ cells per sample were washed with 1% BSA diluted in FLUO-Buffer (1x PBS + 1%BSA + 0.01% NaN3). To discriminate viable cells, the samples were incubated with Fixable Viability Dye eFluor 780 (eBioscience, Thermo Fisher Scientific, USA). Following a wash step with FLUO-Buffer, the cells were incubated with primary antibodies for 20 min, rewashed with FLUO-Buffer, and subsequently incubated with conjugated secondary antibodies for 20 min. After a final wash step, the cells were analyzed using the AttuneNXT flow cytometer (Thermo Fisher Scientific, USA)^73^. Data analysis was performed using FlowJo 10.8.1 software (FlowJo, Ashland, USA). Representative gating strategies employed for the first and second experiments are illustrated in Supplementary Figs. 1 and 5, respectively.

### Microbiome analysis

Cecum content was taken from 4-week-old WT and IFNLR1^−/−^ chickens. The caeca were isolated from the rest of the intestine, and one cecum was cut open longitudinally. The content was collected using sterile 10µl inoculation loops and transferred to 2 ml lysing matrix B tubes (MP Biomedicals) filled with 1 ml DNA Stool Stabilizer (Invitek Diagnostics, Germany). The remaining digesta was cautiously removed and the mucosa was scrapped with the inoculation loop to transfer mucosa-associated bacteria. Samples were kept on ice and brought to the laboratory within 2h after collection. DNA was extracted and processed at the TUM Core Facility Microbiome. In brief, following mechanical cell lysis, DNA was purified on columns (Machery-Nagel, Düren, Germany). The V3/V4 region of the 16S rRNA gene was amplified by 25 cycles of a two-step PCR with primers 341FW and 785RV using a combinatorial dual-barcoding strategy. Amplicon libraries were sequenced on an Illumina MiSeq (Illumina, San Diega, USA) following the manufacturer’s instructions. A detailed description of DNA isolation, library preparation, and sequencing can be found elsewhere^74^. Raw sequence reads were processed using IMNGS2 (www.imngs2.org)^75^, a platform based on UPARSE^76^. Zero-radius operational taxonomic units (zOTUs) were calculated, and a 0.25% abundance filter was applied. The IMNGS2-created output files were further processed using RHEA^77^, a modular pipeline for analyzing microbial profiles based on the 16S rRNA gene. Sequence reads were normalized using the normalization script in RHEA, and sequencing depth was sufficient, as depicted by the rarefaction curve (Supplementary Fig. 2a).

### Enzyme-linked immunosorbent assay (ELISA)

ELISA was performed according to an earlier protocol with minor modifications^69^. For the quantification of total plasma IgM and IgY levels, 96-well maxisorp non-immuno ELISA plates (Thermo Fisher Scientific, Germany) were coated overnight at 4°C with 2 μg/ml goat anti-chicken IgM antibody (Bethyl Laboratories) or 2 μg/ml rabbit anti-chicken IgY antibody (Sigma-Aldrich, Darmstadt, Germany). Subsequently, the plates were blocked with PBS containing 4% skim milk and incubated with chicken plasma samples. Purified IgM (Rockland) at a concentration of 10 μg/ml and IgY (Jackson IR) at 2.9 μg/ml were used as standards. Bound IgM or IgY were detected using goat anti-chicken IgM-horseradish peroxidase (HRP) conjugate (Bethyl Laboratories) or rabbit anti-chicken IgY-HRP conjugate (Sigma-Aldrich, Darmstadt, Germany), respectively, followed by development with 3,3’,5,5’-tetramethylbenzidine (TMB) substrate (Thermo Fisher Scientific, Germany). The OD was measured using a FluoStar Omega microplate reader (BMG LABTECH, Ortenberg, Germany) at a wavelength of 450 nm with a reference filter of 620 nm (Version 5.91 5.70 R2). To determine antigen-specific IgM and IgY levels, the ELISA plates were coated with 2 μg/ml KLH (Sigma-Aldrich, Darmstadt, Germany) or BSA, and subsequent steps were performed as described for the detection of total plasma IgM and IgY, without the inclusion of purified immunoglobulin standards.

### TCR Amplicon Sequencing

T cell receptor (TCR) amplicons were prepared for Next Generation Sequencing (NGS) as previously described^78^. Briefly, total RNA was extracted from the spleen using the SV Total RNA Isolation System (Promega, Madison, USA), quantified on a Nanodrop ND-1000 (Thermo Fisher Scientific, Waltham, USA), and the quality was determined on a 2100 Bioanalyzer (Agilent Technologies, Santa Clara, USA). 900 ng of total RNA from samples with an RNA Integrity Number (RIN) ≥ 8 were subjected to TCR constant (C) gene-specific 5’RACE with molecular barcoding followed by two consecutive semi-nested and semi-step-out PCR reactions according to previously established protocols^78^. The number of repeated cycles was 21 in the first PCR and 11 (β) or 13 (α) in the second PCR. Gel-purified PCR amplicons were sequenced on an Illumina MiSeq Instrument (Illumina, San Diego, USA) by Eurofins Genomics Germany GmbH (Ebersberg, Germany)

### TCR repertoire analysis

Raw read quality assessment with *FastQC v0.12.1*^79^, available online at: https://www.bioinformatics.babraham.ac.uk/projects/fastqc/.) and *MultiQC v1.23*^80^ was followed by TCR analysis and annotation with *MiXCR v4.7.0*^81^ as previously described^78^. CDR3 clonotypes were annotated and quantified using a chicken-specific germline V(D)J gene segment library with *MiXCR* and analyzed with *R v4.4.1*. Samples were down sampled with the smallest sample size greater than 0.5 * 20^th^ quantile as the target size. Analysis was performed using *tidyverse v2.0.0.* packages. For the Kmer analysis, the first and last 4 amino acids of the CDR3 were excluded, and kmer counts were normalized to the total count of CDR3s in the sample.

### RNA isolation, cDNA synthesis, and qRT-PCR

Total RNA was isolated from the CAM and spleen using the Trizol method (VWR, Germany)^68^. The concentration of the extracted RNA was assessed by measuring the absorbance ratio at 260/280 nm using a Biospec-nano Spectrophotometer. An absorbance ratio between 1.9 and 2.0 was considered indicative of pure RNA. Bioanalyzer Systems 6000 nano kit was also used to measure RNA integrity, and only RIN values above 7 were considered for further processing. Subsequently, 400 ng of RNA was reverse transcribed into complementary DNA (cDNA) using the GoScript™ Reverse Transcription Mix and Random Primers (Promega, USA), following the manufacturer’s protocol. The primer sequences used to analyze various gene expressions in the CAM and spleen are provided in Supplementary Table 6. A no-template control (NTC), containing PCR-grade water instead of the sample, was included to detect potential contamination. Real-time quantitative reverse transcription PCR (qRT-PCR) was performed using the QuantStudio5 system (QuantStudio™ Design&analysis Software v1.5.2, Thermo Fisher Scientific) and the Promega GoTaq qRT-PCR kit (Promega, USA), following the manufacturer’s instructions. The annealing temperature for all the studied genes was 59°C. The expression of each gene was analyzed in triplicate and normalized to the 18S ribosomal RNA (rRNA) gene, which served as the housekeeping reference. For the third experiment, the mRNA abundance in unchallenged WT embryos was used to calculate the relative expression levels across all experimental groups. For the spleen analysis of H3N1 infection, the challenged WT group was used as a control. In addition, the genes that were downregulated in IFNAR1^−/−^ were presented as log2. The relative quantification of the studied genes was done using the 2^−ΔΔCt^ method.

### Scoring and histology of the oviduct

Lesions were scored in the reproductive tract in all *in vivo* H3N1-challenged groups and negative controls^30^. Magnum grade 0: normal with physiological height of mucosa, Magnum grade 1: mild lymphoplasmacellular salpingitis with physiological height of mucosa, Magnum grade 2: moderate lymphoplasmacellular salpingitis with prominent atrophy of the mucosa, Magnum grade 3: severe lymphoplasmacellular salpingitis with severe atrophy of the mucosa. The oviduct (infundibulum, magnum) of day 5 post-infection of the unchallenged WT group, as well as WT and IFNLR1^−/−^ (challenged with H3N1) were used for histological assessment. The samples were fixed in 10% neutral buffered formalin and processed routinely. Four-micrometer tissue sections were stained with hematoxylin and eosin for light microscopy. The sections’ analysis addressed histopathological changes and signs of inflammatory reactions, including hyperemia, edema, loss of epithelial cells, and infiltration of immune cells.

### Statistical analysis

Statistical analyses were performed using SPSS statistics software (version 28.0.1.1. IBM). The data obtained from the study were analyzed either with one-way ANOVA or two-way ANOVA based on the experimental design. All p-values ≤ 0.05 were marked as significant. Graphs were designed with GraphPad Prism (Version 9.3.1. Dotmatics). Statistics of the microbiome analysis were performed using the Wilcoxon rank-sum pairwise test and PERMANOVA. For the TCR repertoire, statistical analyses were conducted in *R* as previously described^78^. A negative binomial generalized linear model (GLM) was fitted using the *glm.nb()* function from the *MASS* package v7.3-61^82^ available at: https://www.stats.ox.ac.uk/pub/MASS4/.) with TCR clonotype counts of V genes as the response variable and the experimental group (Genotype and Immunization) as predictors. *Post-hoc* pairwise comparisons among the levels of predictors were conducted with the *emmeans* package v1.10.3^83^, available at: https://socialsciences.mcmaster.ca/jfox/Books/Companion/.) applying Holm adjustment for multiple comparisons.

## Acknowledgments

The TUM Animal Research Center (ARC) provided animal husbandry and experimental infrastructure. The authors are grateful to the ARC animal keepers for their dedicated support. The authors also acknowledge Poulpharm Belgium’s valuable support and contributions to this research by providing the H3N1 virus isolate. ChatGPT (https://chat.openai.com/) was used to improve the conciseness of language, with no contribution to the scientific content.

## Data availability

The data supporting the findings of this study are available in the article and its Supplementary Information section. Further data are available upon reasonable request from the corresponding author.

## Author contributions

Conceptualization: M.N.A., and B. Schusser. Methodology and Investigation: M.N.A., H.V., R.K., C.Z., S.P.F., T.v.H., S.S., M.B., R.N., T.VL.B., L.A., A.R., B.S., B.A.A., S.R., R.P., H.S., and B. Schusser. Visualization: M.A., C.Z., and S.P.F. Funding acquisition: M.N.A., and B. Schusser. Supervision: B. Schusser. Writing-original draft: M.N.A. All authors reviewed and edited the manuscript.

## References

1. Peacock, T.P., James, J., Sealy, J.E. & Iqbal, M. A global perspective on H9N2 avian influenza virus. Viruses 11, 620 (2019).

2. Pinotti, F. et al. Modelling the transmission dynamics of H9N2 avian influenza viruses in a live bird market. Nature Communications 15, 3494 (2024).

3. Wong, S.S. & Yuen, K.-y. Avian influenza virus infections in humans. Chest 129, 156–168 (2006).

4. Organization, W.H. Ongoing avian influenza outbreaks in animals pose risk to humans. Situation Analysis and Advice to Countries from FAO, WHO, WOAH. Available online: https://www.who.int/news/item/12-07-2023-ongoing-avianinfluenza-outbreaks-in-animals-pose-risk-to-humans (accessed on 25 October 2023) (2023).

5. Horman, W.S., Nguyen, T.H., Kedzierska, K., Bean, A.G. & Layton, D.S. The drivers of pathology in zoonotic avian influenza: the interplay between host and pathogen. Frontiers in immunology 9, 1812 (2018).

6. Peacock, T. et al. The global H5N1 influenza panzootic in mammals. Nature, 1–2 (2024).

7. Seeger, R.M., Hagerman, A.D., Johnson, K.K., Pendell, D.L. & Marsh, T.L. When poultry take a sick leave: Response costs for the 2014–2015 highly pathogenic avian influenza epidemic in the USA. Food Policy 102, 102068 (2021).

8. Guinat, C. et al. Promising Effects of Duck Vaccination against Highly Pathogenic Avian Influenza, France, 2023–2024. Emerging Infectious Diseases 31, 1468–1471 (2025).

9. Lindenmann, J., Burke, D. & Isaacs, A. Studies on the production, mode of action and properties of interferon. British journal of experimental pathology 38, 551 (1957).

10. Durbin, R.K., Kotenko, S.V. & Durbin, J.E. Interferon induction and function at the mucosal surface. Immunological reviews 255, 25–39 (2013).

11. Reuter, A. et al. Antiviral activity of lambda interferon in chickens. Journal of virology 88, 2835–2843 (2014).

12. Challagulla, A. et al. Germline engineering of the chicken genome using CRISPR/Cas9 by in vivo transfection of PGCs. Animal Biotechnology 34, 775–784 (2023).

13. Santhakumar, D., Rubbenstroth, D., Martinez-Sobrido, L. & Munir, M. Avian Interferons and Their Antiviral Effectors. Front Immunol 8, 49 (2017).

14. Karpala, A.J. et al. Molecular cloning, expression, and characterization of chicken IFN-λ. journal of interferon & cytokine research 28, 341–350 (2008).

15. Galani, I.E. et al. Interferon-λ mediates non-redundant front-line antiviral protection against influenza virus infection without compromising host fitness. Immunity 46, 875–890. e876 (2017).

16. Davidson, S. et al. IFN λ is a potent anti-influenza therapeutic without the inflammatory side effects of IFN α treatment. EMBO molecular medicine 8, 1099–1112 (2016).

17. Lozhkov, A.A. et al. The key roles of interferon lambda in human molecular defense against respiratory viral infections. Pathogens 9, 989 (2020).

18. Mordstein, M. et al. Interferon-λ contributes to innate immunity of mice against influenza A virus but not against hepatotropic viruses. PLoS pathogens 4, e1000151 (2008).

19. Mordstein, M. et al. Lambda interferon renders epithelial cells of the respiratory and gastrointestinal tracts resistant to viral infections. Journal of virology 84, 5670–5677 (2010).

20. Broggi, A. et al. Type III interferons disrupt the lung epithelial barrier upon viral recognition. Science 369, 706–712 (2020).

21. Major, J. et al. Type I and III interferons disrupt lung epithelial repair during recovery from viral infection. Science 369, 712–717 (2020).

22. Penski, N. et al. Highly pathogenic avian influenza viruses do not inhibit interferon synthesis in infected chickens but can override the interferon-induced antiviral state. Journal of virology 85, 7730–7741 (2011).

23. Schoggins, J.W. Interferon-stimulated genes: roles in viral pathogenesis. Curr Opin Virol 6, 40–46 (2014).

24. McNab, F., Mayer-Barber, K., Sher, A., Wack, A. & O’garra, A. Type I interferons in infectious disease. Nature Reviews Immunology 15, 87–103 (2015).

25. Caine, E.A. et al. Interferon lambda protects the female reproductive tract against Zika virus infection. Nature communications 10, 280 (2019).

26. Cumming, H.E. & Bourke, N.M. Type I IFNs in the female reproductive tract: The first line of defense in an ever-changing battleground. Journal of leukocyte biology 105, 353–361 (2019).

27. Ank, N. et al. An important role for type III interferon (IFN-lambda/IL-28) in TLR-induced antiviral activity. J Immunol 180, 2474–2485 (2008).

28. Müller, U. et al. Functional role of type I and type II interferons in antiviral defense. Science 264, 1918–1921 (1994).

29. de Wit, J. et al. Major difference in clinical outcome and replication of a H3N1 avian influenza strain in young pullets and adult layers. Avian Pathology 49, 286–295 (2020).

30. Sid, H. et al. Reinstatement of RIG-I in chickens via genetic modification reveals new insights into the dynamic evolution of avian immune sensors. Frontiers in immunology 16, (2025).

31. Kaiser, P. Advances in avian immunology—prospects for disease control: a review. Avian pathology 39, 309–324 (2010).

32. Bean, A.G. & Lowenthal, J.W. Avian cytokines and their receptors. Avian immunology. Elsevier, 2022, pp 249–276.

33. Eyerich, S. et al. Th22 cells represent a distinct human T cell subset involved in epidermal immunity and remodeling. The Journal of clinical investigation 119, 3573–3585 (2009).

34. Dudakov, J.A., Hanash, A.M. & van den Brink, M.R. Interleukin-22: immunobiology and pathology. Annual review of immunology 33, 747–785 (2015).

35. Katze, M.G., He, Y. & Gale, M. Viruses and interferon: a fight for supremacy. Nature Reviews Immunology 2, 675–687 (2002).

36. Röll, S., Härtle, S., Lütteke, T., Kaspers, B. & Härtle, S. Tissue and time specific expression pattern of interferon regulated genes in the chicken. BMC Genomics 18, 264 (2017).

37. Galani, I.-E. et al. Untuned antiviral immunity in COVID-19 revealed by temporal type I/III interferon patterns and flu comparison. Nature immunology 22, 32–40 (2021).

38. Hemann, E.A. et al. Interferon-λ modulates dendritic cells to facilitate T cell immunity during infection with influenza A virus. Nature immunology 20, 1035–1045 (2019).

39. Santer, D.M. et al. Interferon-λ treatment accelerates SARS-CoV-2 clearance despite age-related delays in the induction of T cell immunity. Nat Commun 13, 6992 (2022).

40. Bracci, L. et al. Type I IFN is a powerful mucosal adjuvant for a selective intranasal vaccination against influenza virus in mice and affects antigen capture at mucosal level. Vaccine 23, 2994–3004 (2005).

41. Ye, L. et al. Interferon-λ improves the efficacy of intranasally or rectally administered influenza subunit vaccines by a thymic stromal lymphopoietin-dependent mechanism. Frontiers in Immunology 12, 749325 (2021).

42. Wu, Z. & Kaiser, P. Antigen presenting cells in a non-mammalian model system, the chicken. Immunobiology 216, 1177–1183 (2011).

43. Silva, A.P.d. & Gallardo, R.A. The chicken MHC: insights into genetic resistance, immunity, and inflammation following infectious bronchitis virus infections. Vaccines 8, 637 (2020).

44. Ashby, K.M. et al. Sterile production of interferons in the thymus affects T cell repertoire selection. Science Immunology 9, eadp1139 (2024).

45. Chen, H. et al. Clinical and epidemiological characteristics of a fatal case of avian influenza A H10N8 virus infection: a descriptive study. The Lancet 383, 714–721 (2014).

46. Pu, J. et al. Evolution of the H9N2 influenza genotype that facilitated the genesis of the novel H7N9 virus. Proceedings of the National Academy of Sciences 112, 548–553 (2015).

47. Nowak-Sliwinska, P., Segura, T. & Iruela-Arispe, M.L. The chicken chorioallantoic membrane model in biology, medicine and bioengineering. Angiogenesis 17, 779–804 (2014).

48. Raj, G.D. & Jones, R. Local antibody production in the oviduct and gut of hens infected with a variant strain of infectious bronchitis virus. Veterinary immunology and immunopathology 53, 147–161 (1996).

49. Yang, W., Liu, X. & Wang, X. The immune system of chicken and its response to H9N2 avian influenza virus. Veterinary Quarterly 43, 1–14 (2023).

50. Swayne, D.E. & Pantin-Jackwood, M. Pathobiology of avian influenza virus infections in birds and mammals. Avian influenza 28, 87–122 (2008).

51. Schön, J. et al. Neuraminidase-associated plasminogen recruitment enables systemic spread of natural avian Influenza viruses H3N1. PLoS Pathog 17, e1009490 (2021).

52. Mork, A.-K. et al. Differences in the tissue tropism to chicken oviduct epithelial cells between avian coronavirus IBV strains QX and B1648 are not related to the sialic acid binding properties of their spike proteins. Veterinary research 45, 1–10 (2014).

53. Goel, R.R., Kotenko, S.V. & Kaplan, M.J. Interferon lambda in inflammation and autoimmune rheumatic diseases. Nature Reviews Rheumatology 17, 349–362 (2021).

54. Qu, C.K. The SHP-2 tyrosine phosphatase: signaling mechanisms and biological functions. Cell research 10, 279–288 (2000).

55. Sheehan, K.C., Lazear, H.M., Diamond, M.S. & Schreiber, R.D. Selective blockade of interferon-α and-β reveals their non-redundant functions in a mouse model of West Nile virus infection. PloS one 10, e0128636 (2015).

56. Seo, S.-U. et al. Type I interferon signaling regulates Ly6Chi monocytes and neutrophils during acute viral pneumonia in mice. PLoS pathogens 7, e1001304 (2011).

57. Sharma, L. et al. Distinct roles of type I and type III interferons during a native murine β coronavirus lung infection. Journal of virology 96, e01241–01221 (2022).

58. Burkhardt, N.B. et al. The discovery of chicken foxp3 demands redefinition of avian regulatory T cells. The Journal of Immunology 208, 1128–1138 (2022).

59. Herold, S. et al. Lung epithelial apoptosis in influenza virus pneumonia: the role of macrophage-expressed TNF-related apoptosis-inducing ligand. The Journal of experimental medicine 205, 3065–3077 (2008).

60. Lin, K.L., Suzuki, Y., Nakano, H., Ramsburg, E. & Gunn, M.D. CCR2+ monocyte-derived dendritic cells and exudate macrophages produce influenza-induced pulmonary immune pathology and mortality. The Journal of Immunology 180, 2562–2572 (2008).

61. Hamburger, V. & Hamilton, H.L. A series of normal stages in the development of the chick embryo. Journal of morphology 88, 49–92 (1951).

62. Whyte, J. et al. FGF, insulin, and SMAD signaling cooperate for avian primordial germ cell self-renewal. Stem cell reports 5, 1171–1182 (2015).

63. Hellmich, R. et al. Acquiring resistance against a retroviral infection via CRISPR/Cas9 targeted genome editing in a commercial chicken line. Frontiers in Genome Editing 2, 3 (2020).

64. Schusser, B. et al. Immunoglobulin knockout chickens via efficient homologous recombination in primordial germ cells. Proc Natl Acad Sci U S A 110, 20170–20175 (2013).

65. Van de Lavoir, M.-C. et al. Germline transmission of genetically modified primordial germ cells. Nature 441, 766–769 (2006).

66. Hernandez, R. & Brown, D.T. Growth and maintenance of chick embryo fibroblasts (CEF). Current protocols in microbiology 17, A. 4I. 1–A. 4I. 8 (2010).

67. Schusser, B. et al. Mx is dispensable for interferon-mediated resistance of chicken cells against influenza A virus. Journal of virology 85, 8307–8315 (2011).

68. Panda, B.S. et al. Proteomics and transcriptomics study reveals the utility of ISGs as novel molecules for early pregnancy diagnosis in dairy cows. Journal of Reproductive Immunology 140, 103148 (2020).

69. Schusser, B. et al. Expression of heavy chain-only antibodies can support B-cell development in light chain knockout chickens. Eur J Immunol 46, 2137–2148 (2016).

70. Brauer, R. & Chen, P. Influenza virus propagation in embryonated chicken eggs. Journal of visualized experiments: JoVE (2015).

71. Schultz, U., Rinderle, C., Sekellick, M.J., Marcus, P.I. & Staeheli, P. Recombinant chicken interferon from Escherichia coli and transfected COS cells is biologically active. Eur J Biochem 229, 73–76 (1995).

72. Sid, H., Hartmann, S., Winter, C. & Rautenschlein, S. Interaction of influenza A viruses with oviduct explants of different avian species. Frontiers in Microbiology 8, 1338 (2017).

73. Cossarizza, A. et al. Guidelines for the use of flow cytometry and cell sorting in immunological studies. European journal of immunology 51, 2708–3145 (2021).

74. Reitmeier, S., Kiessling, S., Neuhaus, K. & Haller, D. Comparing circadian rhythmicity in the human gut microbiome. STAR protocols 1, 100148 (2020).

75. Lagkouvardos, I. et al. IMNGS: a comprehensive open resource of processed 16S rRNA microbial profiles for ecology and diversity studies. Scientific reports 6, 33721 (2016).

76. Edgar, R.C. UPARSE: highly accurate OTU sequences from microbial amplicon reads. Nature methods 10, 996–998 (2013).

77. Lagkouvardos, I., Fischer, S., Kumar, N. & Clavel, T. Rhea: a transparent and modular R pipeline for microbial profiling based on 16S rRNA gene amplicons. PeerJ 5, e2836 (2017).

78. Früh, S.P., Früh, M.A., Kaufer, B.B. & Göbel, T.W. Unraveling the chicken T cell repertoire with enhanced genome annotation. Frontiers in Immunology 15, 1359169 (2024).

79. Andrews, S. FastQC: a quality control tool for high throughput sequence data. Cambridge, United Kingdom; 2010.

80. Ewels, P., Magnusson, M., Lundin, S. & Käller, M. MultiQC: summarize analysis results for multiple tools and samples in a single report. Bioinformatics 32, 3047–3048 (2016).

81. Bolotin, D.A. et al. MiXCR: software for comprehensive adaptive immunity profiling. Nature methods 12, 380–381 (2015).

82. Venables, W. & Ripley, B. Modern applied statistics with S fourth. New York: Springer. (2002).

83. Fox, J. & Weisberg, S. An R companion to applied regression. Sage publications, 2018.

