## Supplementary Information for "Deciphering interferon functions in avian influenza: Insights from receptor knockout models in the natural host"

### Supplementary materials for

#### Type I and type III interferon receptor knockout chickens: Novel models for unraveling interferon dynamics in influenza infection

Mohanned Naif Alhussien[1], Hanna-Kaisa Vikkula[1], Romina Klinger[1], Christian Zenner[1], Simon P Früh[2,3], Rashi Negi[1], Theresa von Heyl[1], Sabrina Schleibinger[1], Milena Brunner[1], Tom VL Berghof[1], Leora Avolio[1], Arne Reich[1], Benjamin Schade[4], Bassel A. Abukhadra[5], Silke Rautenschlein[5], Rudolf Preisinger[6], Hicham Sid[1], Benjamin Schusser[1,7*]

[1]TUM School of Life Sciences, Weihenstephan, Department of Molecular Life Sciences, Reproductive Biotechnology, Technical University of Munich, 85354, Freising, Germany
[2]Department of Veterinary Sciences, Ludwig-Maximilians-Universität München, Munich, Germany.
[3]Institute of Virology, Freie Universität Berlin, Berlin, Germany.
[4]Bavarian Animal Health Service, Department of Pathology, 85586, Poing, Germany
[5]Clinic for Poultry, University of Veterinary Medicine Hannover, Hannover, Germany
[6]EW GROUP GmbH, Visbek, Germany
[7]Center for Infection Prevention (ZIP), Technical University of Munich, 85354 Freising, Germany

This document contains Supplementary Figures 1-12 and Supplementary Tables 1-6

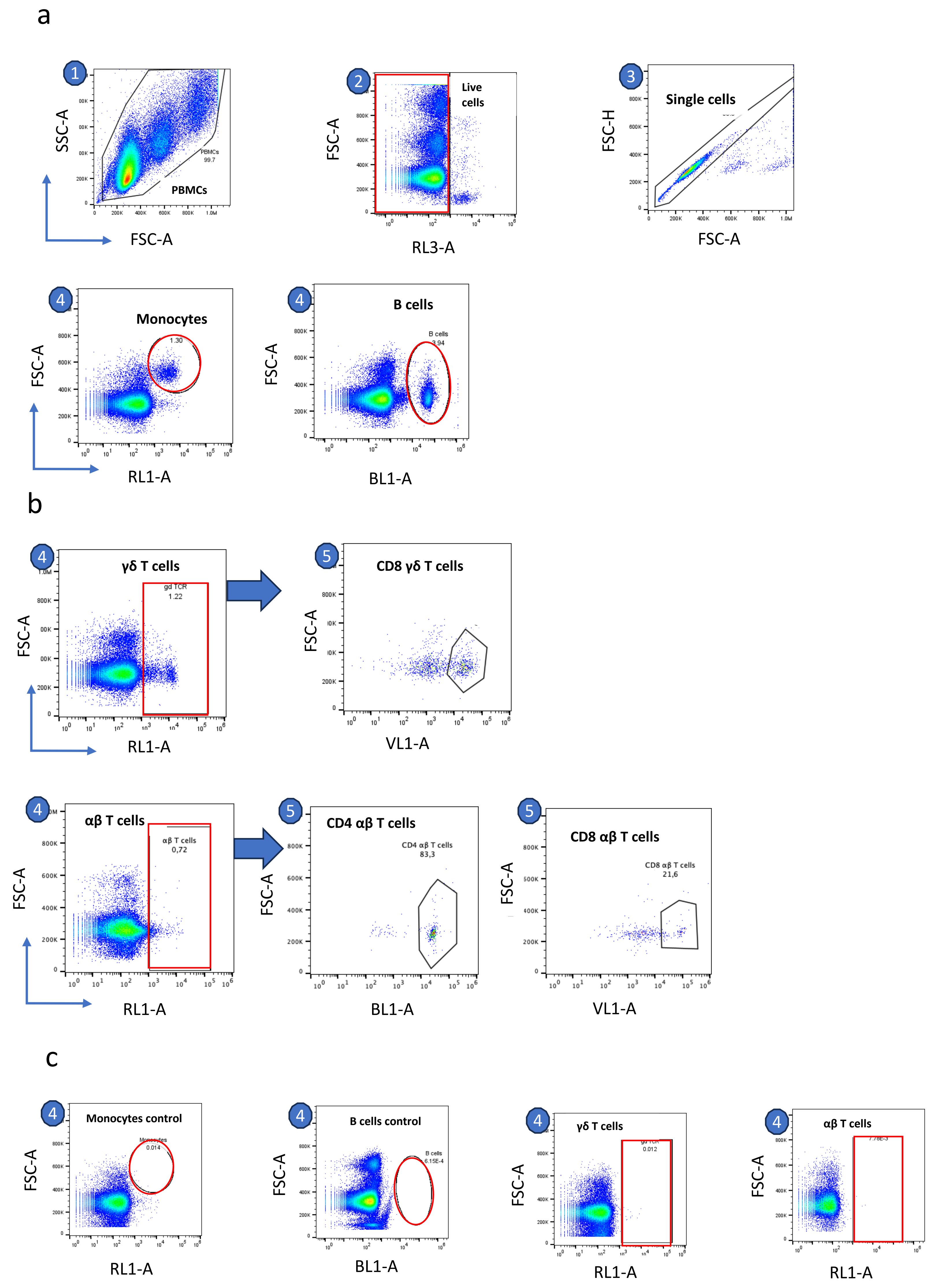
a
1
SSC-A
FSC-A
PBMCs
PBMCs
99.7
2
Live cells
FSC-A
RL3-A
3
Single cells
FSC-H
FSC-A
4
Monocytes
1.30
FSC-A
RL1-A
4
B cells
B cells
3.94
FSC-A
BL1-A
b
4
γδ T cells
gd TCR
1.22
FSC-A
RL1-A
5
CD8 γδ T cells
FSC-A
VL1-A
4
αβ T cells
αβ T cells
0,72
FSC-A
RL1-A
5
CD4 αβ T cells
CD4 αβ T cells
83,3
FSC-A
BL1-A
5
CD8 αβ T cells
CD8 αβ T cells
21,6
FSC-A
VL1-A
c
4
Monocytes control
Monocytes
0.014
FSC-A
RL1-A
4
B cells control
B cells
6.15E-4
FSC-A
BL1-A
4
γδ T cells
0.012
FSC-A
RL1-A
4
αβ T cells
FSC-A
RL1-A

**Supp Fig. 1**

**Supplementary Fig. 1.** Representation of the gating strategy to quantify populations of PBMCs isolated from 1-month-old chicks. Steps 1, 2 & 3 are common for all. All individual cell analyses were gated from step 3 except for the subpopulation of γδ and αβ T cells, which were gated from step 4. **a**, Detection of monocytes and B cells. **b**, detection of γδ, αβ T cells, and their CD4 and CD8 subpopulation. **c**, secondary antibody control used for this experiment.

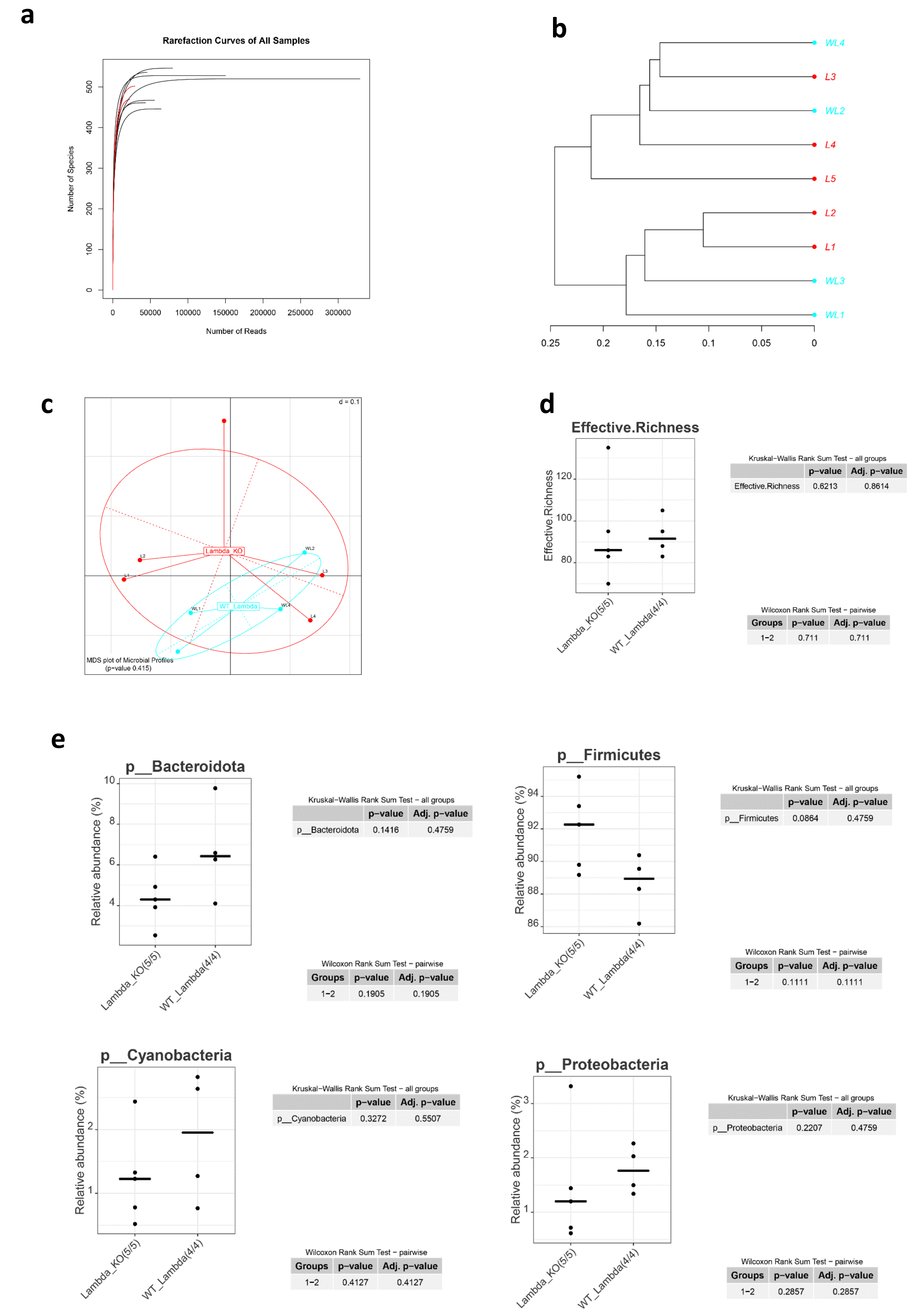
a
Rarefaction Curves of All Samples
Number of Species
Number of Reads
0
100
200
300
400
500
0
50000
100000
150000
200000
250000
300000
b
WL4
L3
WL2
L4
L5
L2
L1
WL3
WL1
0.25
0.2
0.15
0.1
0.05
0
c
d = 0.1
Lambda_KO
WT_Lambda
L1
L2
L3
L4
WL1
WL2
WL4
MDS plot of Microbial Profiles
(p-value 0.415)
d
Effective.Richness
80
100
120
Lambda_KO(5/5)
WT_Lambda(4/4)
Kruskal−Wallis Rank Sum Test − all groups
p−value Adj. p−value
Effective.Richness 0.6213 0.8614
Wilcoxon Rank Sum Test − pairwise
Groups p−value Adj. p−value
1−2 0.711 0.711
e
p__Bacteroidota
Relative abundance (%)
4
6
8
10
Lambda_KO(5/5)
WT_Lambda(4/4)
Kruskal−Wallis Rank Sum Test − all groups
p−value Adj. p−value
p__Bacteroidota 0.1416 0.4759
Wilcoxon Rank Sum Test − pairwise
Groups p−value Adj. p−value
1−2 0.1905 0.1905
p__Firmicutes
Relative abundance (%)
86
88
90
92
94
Lambda_KO(5/5)
WT_Lambda(4/4)
Kruskal−Wallis Rank Sum Test − all groups
p−value Adj. p−value
p__Firmicutes 0.0864 0.4759
Wilcoxon Rank Sum Test − pairwise
Groups p−value Adj. p−value
1−2 0.1111 0.1111
p__Cyanobacteria
Relative abundance (%)
1
2
Lambda_KO(5/5)
WT_Lambda(4/4)
Kruskal−Wallis Rank Sum Test − all groups
p−value Adj. p−value
p__Cyanobacteria 0.3272 0.5507
Wilcoxon Rank Sum Test − pairwise
Groups p−value Adj. p−value
1−2 0.4127 0.4127
p__Proteobacteria
Relative abundance (%)
1
2
3
Lambda_KO(5/5)
WT_Lambda(4/4)
Kruskal−Wallis Rank Sum Test − all groups
p−value Adj. p−value
p__Proteobacteria 0.2207 0.4759
Wilcoxon Rank Sum Test − pairwise
Groups p−value Adj. p−value
1−2 0.2857 0.2857

**Supp Fig. 2**

**Supplementary Fig. 2**. Microbiome analysis of the cecal contents from one-month-old IFNLR1$^{-/-}$ chicks and their WT siblings **a**, Rarefaction curves of all samples show that sequencing depth was sufficient. **b**, A phylogram based on Ward's minimum variance method shows the hierarchical clustering of samples. The sample distance is shown on the x-axis. The IFNLR1$^{-/-}$ chicks are abbreviated as L, and their WT siblings as WL. **c**, Multidimensional scaling plots of microbial profiles. Groups were compared pairwise. The scale bar indicates the distance between samples (d=0.1 describes 10%). **d**, richness as an index for α-diversity. Means are displayed as bold bars. **e**, Taxonomic differences between the groups at the phylum level. Means are indicated as bold bars. The number at the bottom shows the number of observations in the groups.

#### Experiment 2

##### Immunization of IFNAR1 and IFNLR1 knockout chicks

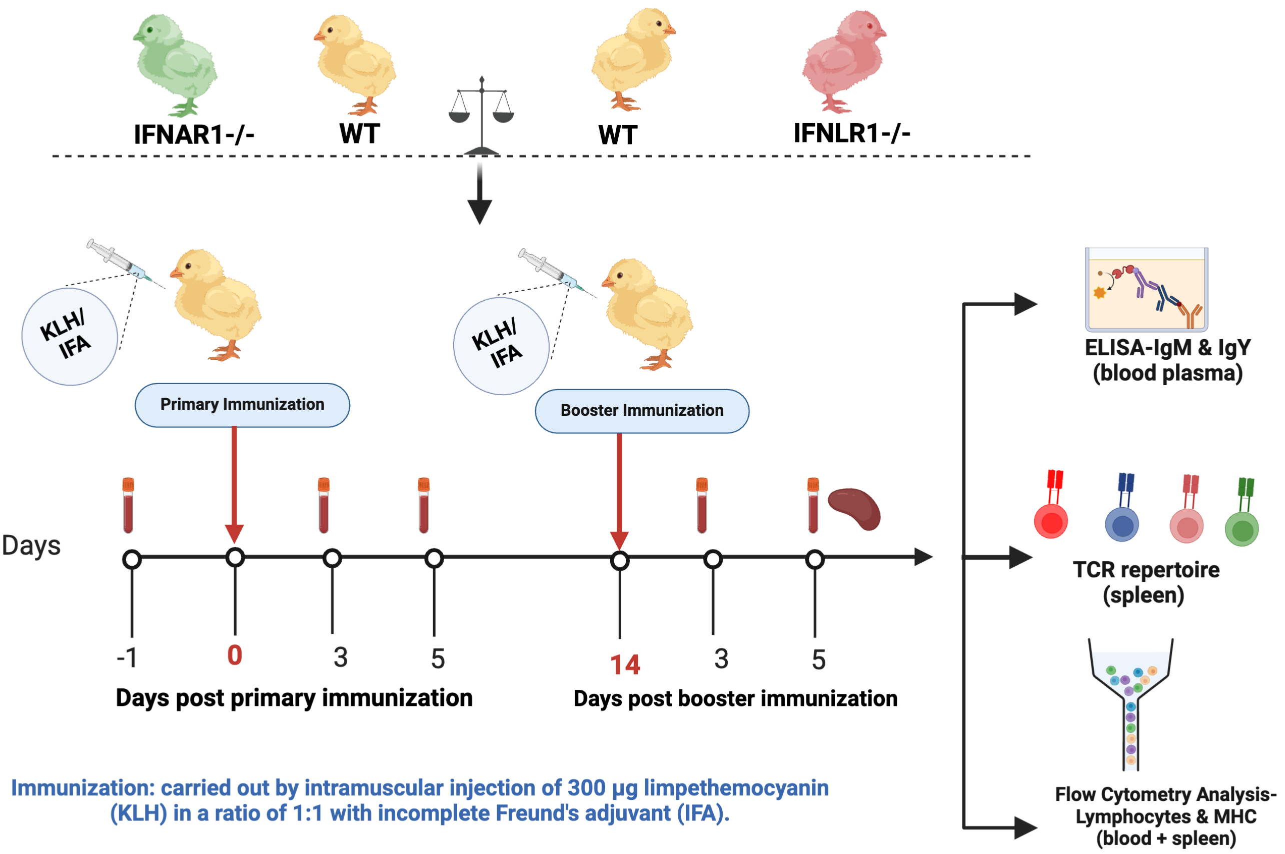


**Supp Fig. 3**

**Supplementary Fig. 3.** In experiment 2, five-week-old WT, IFNLR1$^{-/-}$, and IFNAR1$^{-/-}$ chicks were immunized via intramuscular injection with 300 µg KLH, mixed 1:1 with incomplete Freund's adjuvant. A booster injection of 300 µg KLH, mixed 1:1 with Freund's incomplete adjuvant, was administered two weeks after the initial immunization. Plasma samples were collected from all groups before immunization (day 0), on day 5 post-primary immunization (PP), and on days 3 and 5 post-booster immunization (PB). The total concentration of IgM, IgY, and levels of KLH antigen-specific IgM and IgY were assessed using ELISA. PBMCs were isolated on day 3 post-PP, and splenocytes were isolated on day 5 PB for FACS analysis of immune and MHC-positive cells. T cells of the spleen were used to study TCR repertoire. At least 6 animals per group were included.

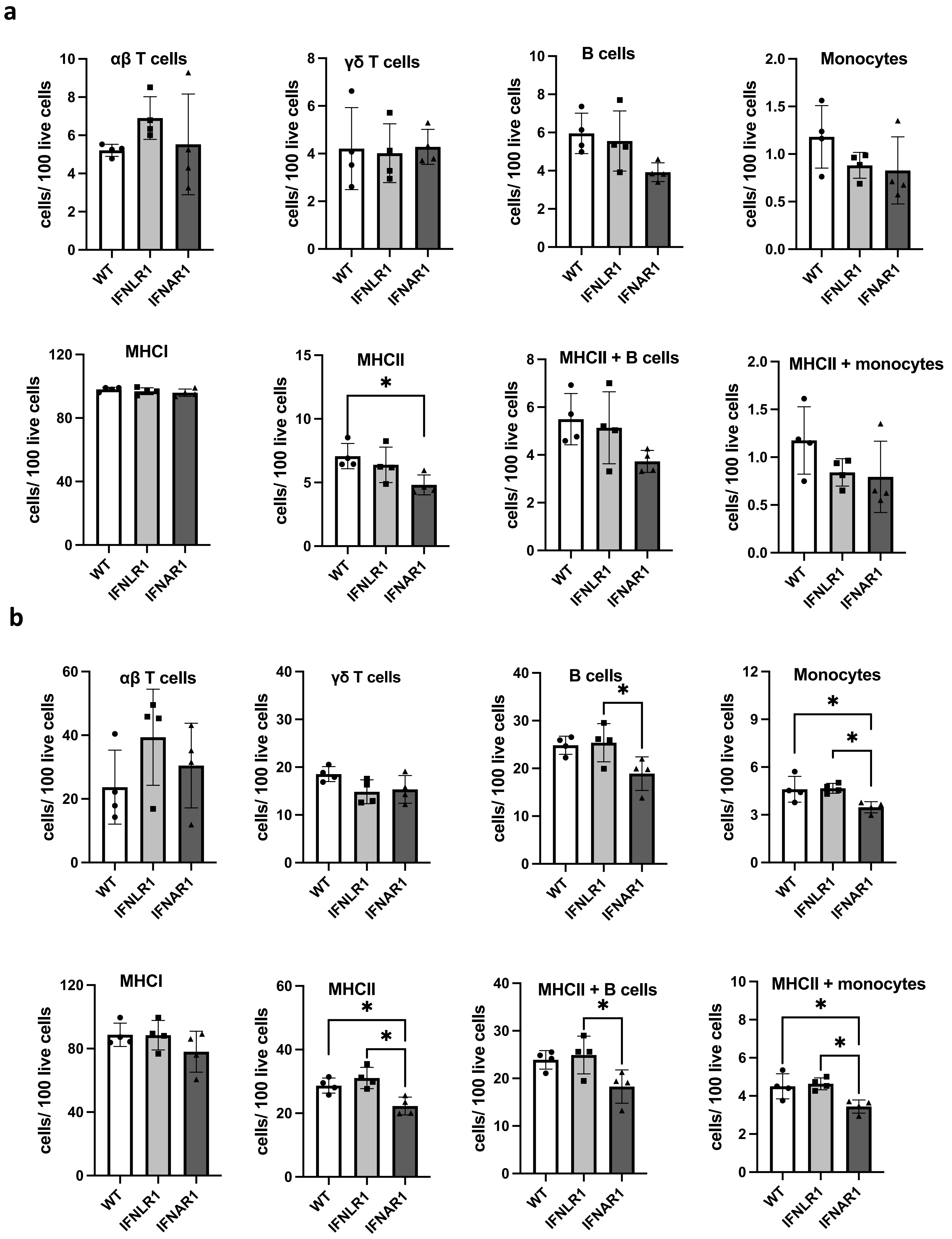
a
αβ T cells
γδ T cells
B cells
Monocytes
MHCI
MHCII
MHCII + B cells
MHCII + monocytes
cells/ 100 live cells
WT
IFNLR1
IFNAR1
b
αβ T cells
γδ T cells
B cells
Monocytes
MHCI
MHCII
MHCII + B cells
MHCII + monocytes

Supp Fig. 4

**Supplementary Fig. 4. Flow cytometry analysis of immune cells and their MHC-positive subpopulations in the blood and spleen of WT, IFNLR1$^{-/-}$ and IFNAR1$^{-/-}$ chicks immunized with KLH. a**, PBMCs were isolated at day 3 post primary immunization and analyzed for differential immune cell populations, including monocytes, B cells, αβ TCR2,3+ or γδ TCR1 + T cells, MHCI, MHCII, MHCII+ B cells, and MHCII+ monocytes. **b,** Splenocytes were isolated at day 5 post booster immunization and analyzed for differential immune cell populations and associated MHC subpopulations. One-way ANOVA was used to analyze the difference between the groups (*****, $P \leq 0.05$).

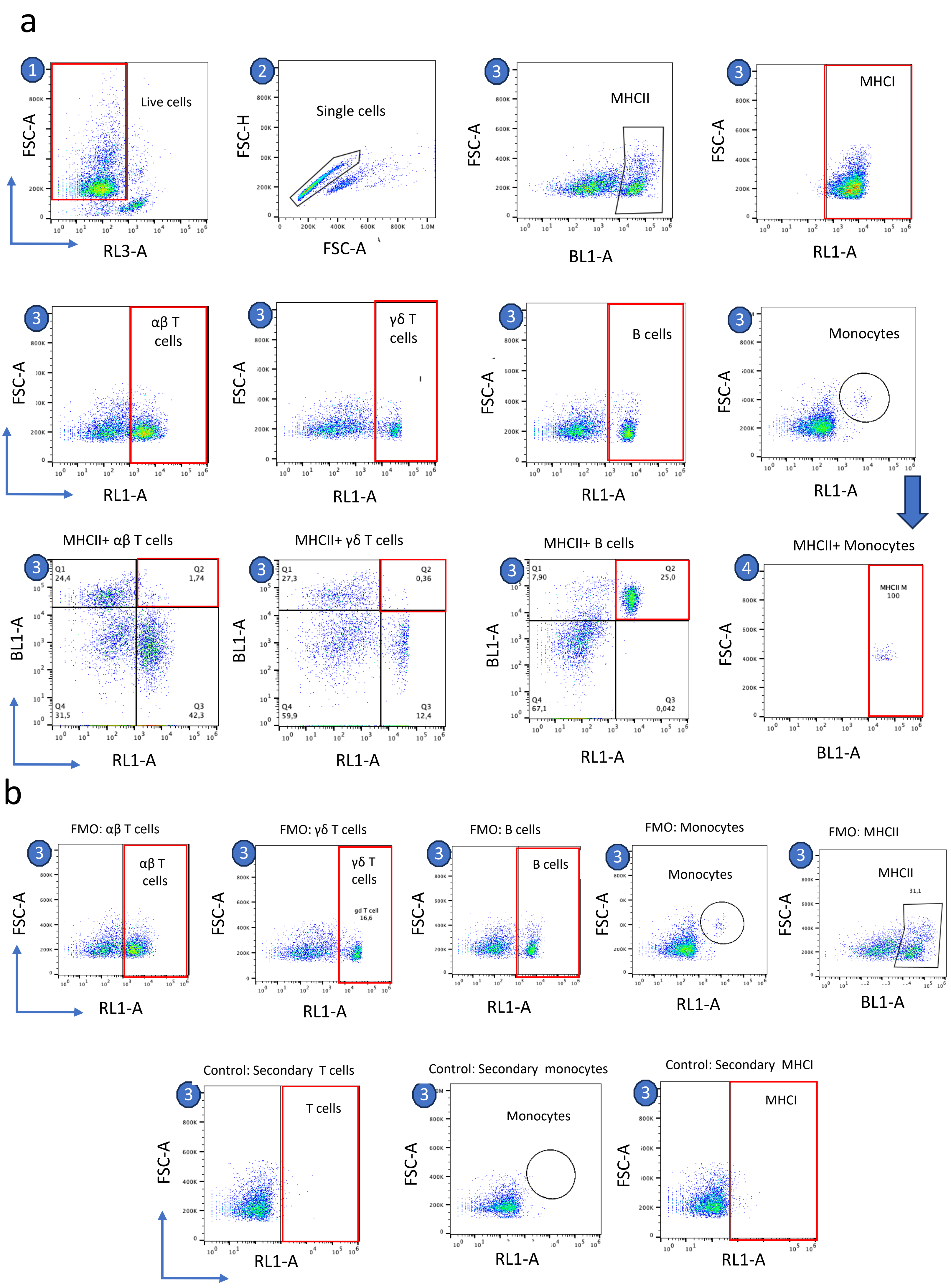
a
Live cells
FSC-A
RL3-A
Single cells
FSC-H
FSC-A
MHCII
BL1-A
MHCI
RL1-A
αβ T cells
γδ T cells
B cells
Monocytes
MHCII+ αβ T cells
Q1 24,4
Q2 1,74
Q4 31,5
Q3 42,3
MHCII+ γδ T cells
Q1 27,3
Q2 0,36
Q4 59,9
Q3 12,4
MHCII+ B cells
Q1 7,90
Q2 25,0
Q4 67,1
Q3 0,042
MHCII+ Monocytes
MHCII M 100
b
FMO: αβ T cells
FMO: γδ T cells
gd T cell 16,6
FMO: B cells
FMO: Monocytes
FMO: MHCII
31,1
Control: Secondary T cells
T cells
Control: Secondary monocytes
Control: Secondary MHCI

**Supp Fig. 5**

**Supplementary Fig. 5.** Representation of gating strategy quantifying immune cells and their MHC-positive subpopulations. Steps 1 and 2 are common for all. All individual cell analyses were gated from step 2 except for MHCII+ monocytes. **a**, immune cells and their subpopulation positive for MHCII. **b**, Fluorescence Minus One (FMO) controls and secondary antibodies control were used for this experiment.

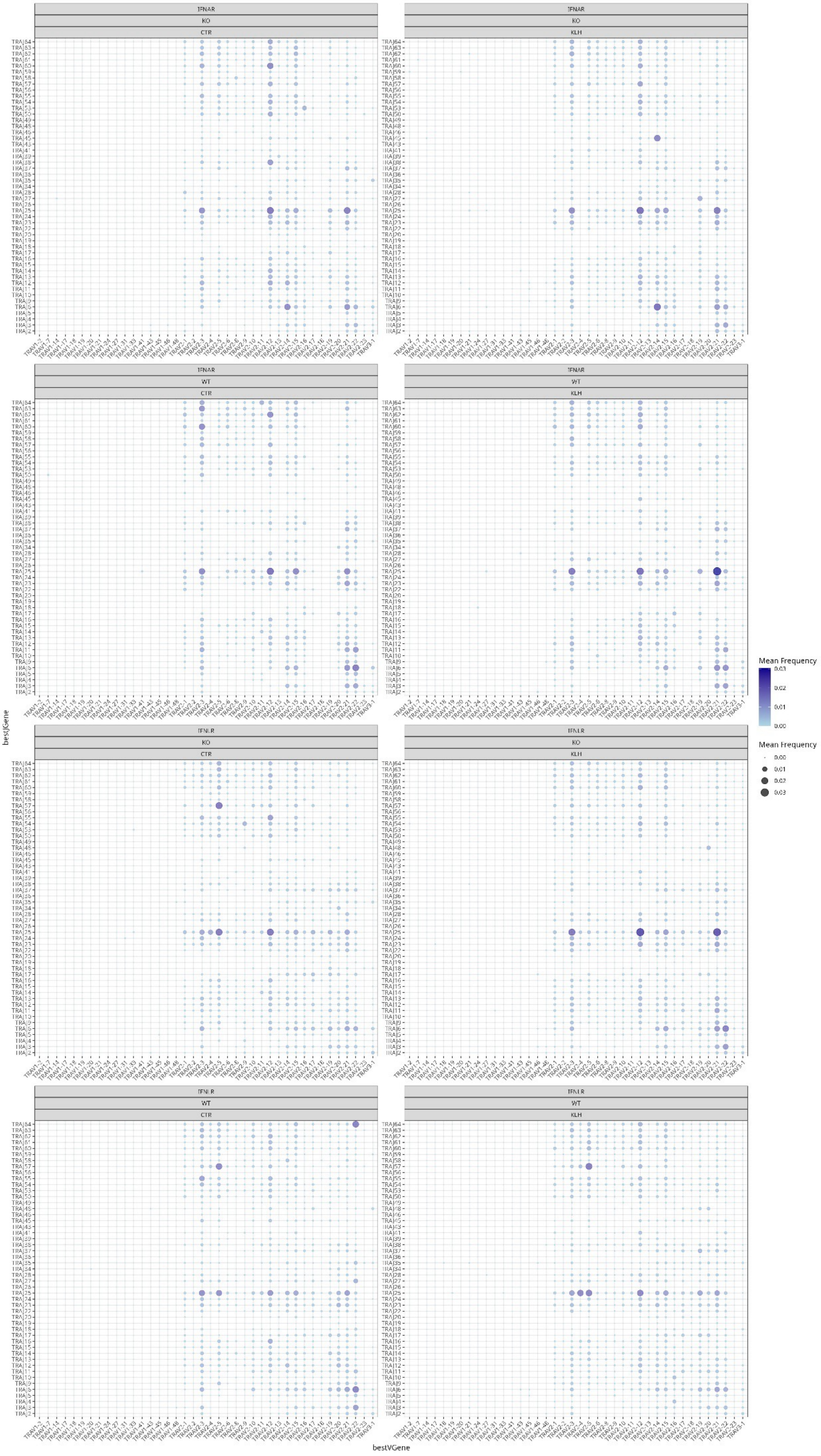
IFNAR
KO
CTR
IFNAR
KO
KLH
IFNAR
WT
CTR
IFNAR
WT
KLH
IFNLR
KO
CTR
IFNLR
KO
KLH
IFNLR
WT
CTR
IFNLR
WT
KLH
bestJGene
bestVGene
Mean Frequency
0.03
0.02
0.01
0.00
Mean Frequency
0.00
0.01
0.02
0.03

**Supp Fig. 6**

**Supplementary Fig. 6. Relative frequency of Vα - Jα genes.**
Mean V-J pairing in TCRα CDR3s in IFNAR1$^{-/-}$, IFNLR1$^{-/-}$, and WT animals in the steady state (CTR) or at day 5 post-KLH booster immunization.

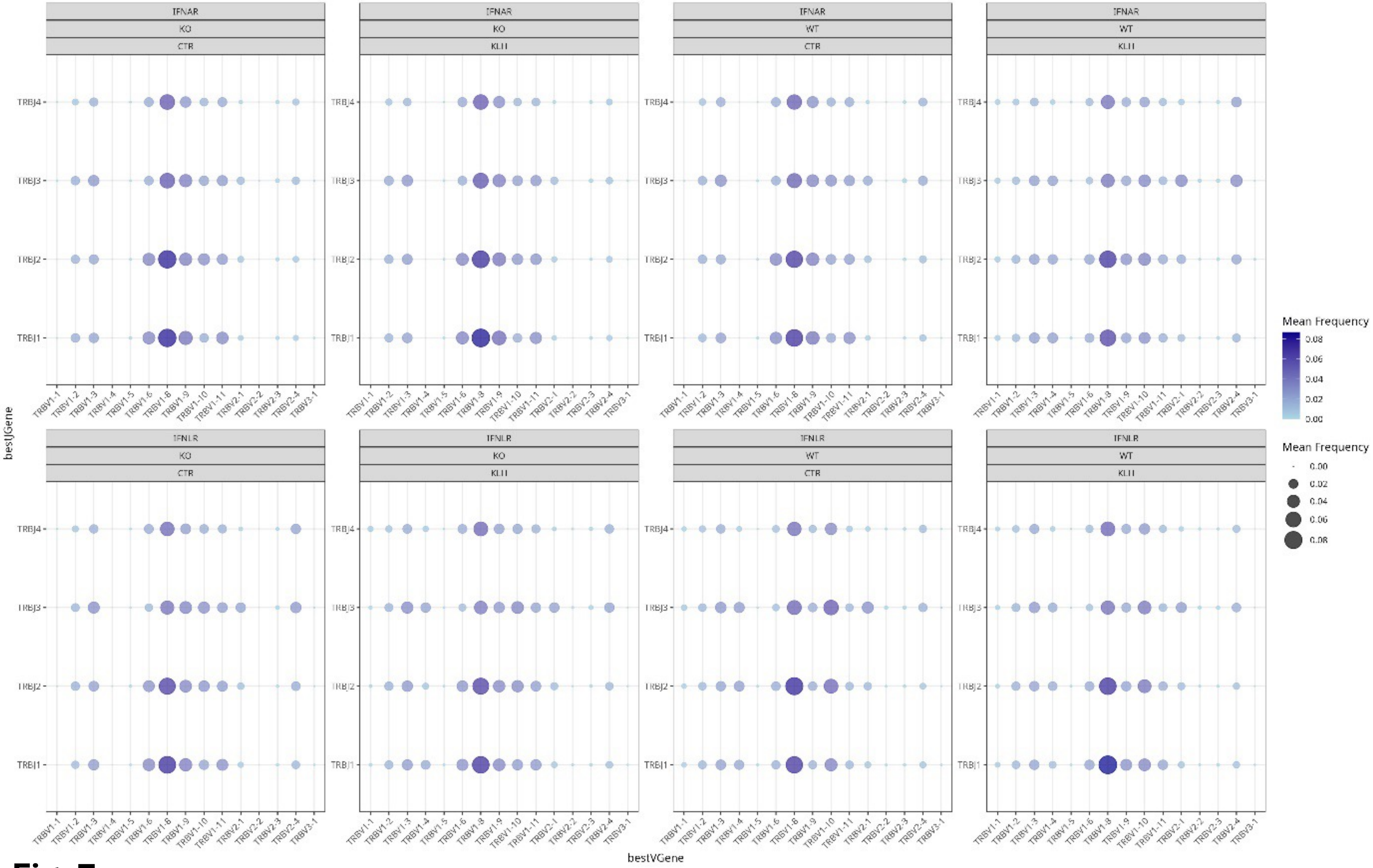
IFNAR
IFNLR
KO
WT
CTR
KLH
TRBJ4
TRBJ3
TRBJ2
TRBJ1
TRBV1-1
TRBV1-2
TRBV1-3
TRBV1-4
TRBV1-5
TRBV1-6
TRBV1-8
TRBV1-9
TRBV1-10
TRBV1-11
TRBV2-1
TRBV2-2
TRBV2-3
TRBV2-4
TRBV3-1
bestJGene
bestVGene
Mean Frequency
0.08
0.06
0.04
0.02
0.00
Mean Frequency
0.00
0.02
0.04
0.06
0.08

**Supp Fig. 7**

**Supplementary Fig. 7. Relative frequency of Vβ - Jβ genes.**
Mean V-J pairing in TCRβ CDR3s in IFNAR1$^{-/-}$, IFNLR1$^{-/-}$, and WT animals in the steady state (CTR) or at day 5 post-KLH booster immunization.

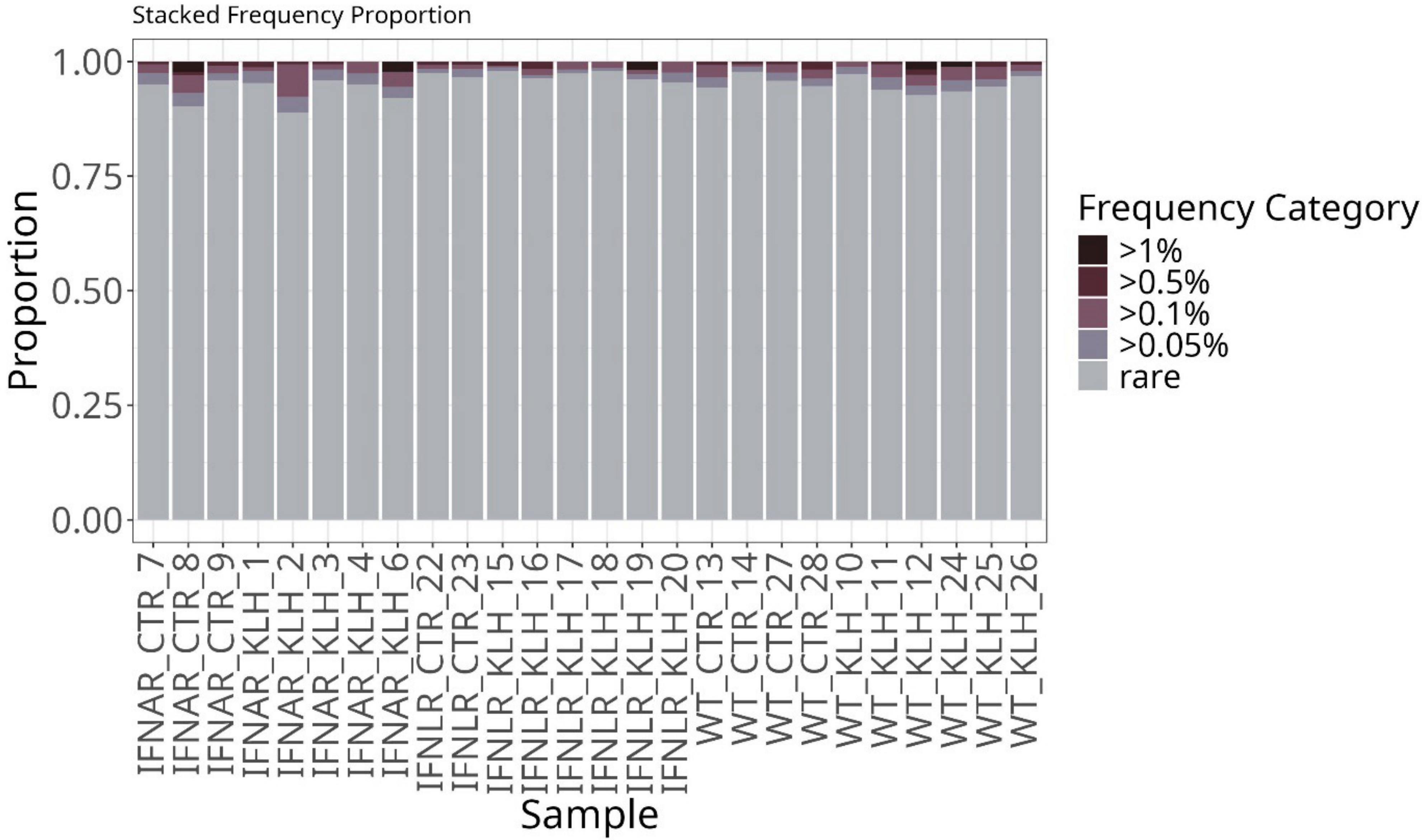
Stacked Frequency Proportion
Proportion
1.00
0.75
0.50
0.25
0.00
IFNAR_CTR_7
IFNAR_CTR_8
IFNAR_CTR_9
IFNAR_KLH_1
IFNAR_KLH_2
IFNAR_KLH_3
IFNAR_KLH_4
IFNAR_KLH_6
IFNLR_CTR_22
IFNLR_CTR_23
IFNLR_KLH_15
IFNLR_KLH_16
IFNLR_KLH_17
IFNLR_KLH_18
IFNLR_KLH_19
IFNLR_KLH_20
WT_CTR_13
WT_CTR_14
WT_CTR_27
WT_CTR_28
WT_KLH_10
WT_KLH_11
WT_KLH_12
WT_KLH_24
WT_KLH_25
WT_KLH_26
Sample
Frequency Category
>1%
>0.5%
>0.1%
>0.05%
rare

**Supp Fig. 8**

**Supplementary Fig. 8**. **Relative frequency distribution of TCRβ CDR3s.**
Distribution of CDR3 clonotypes colored coded by their relative frequencies in IFNAR1$^{-/-}$, IFNLR1$^{-/-}$, and WT animals in the steady state (CTR) or at day 5 post-KLH booster immunization.

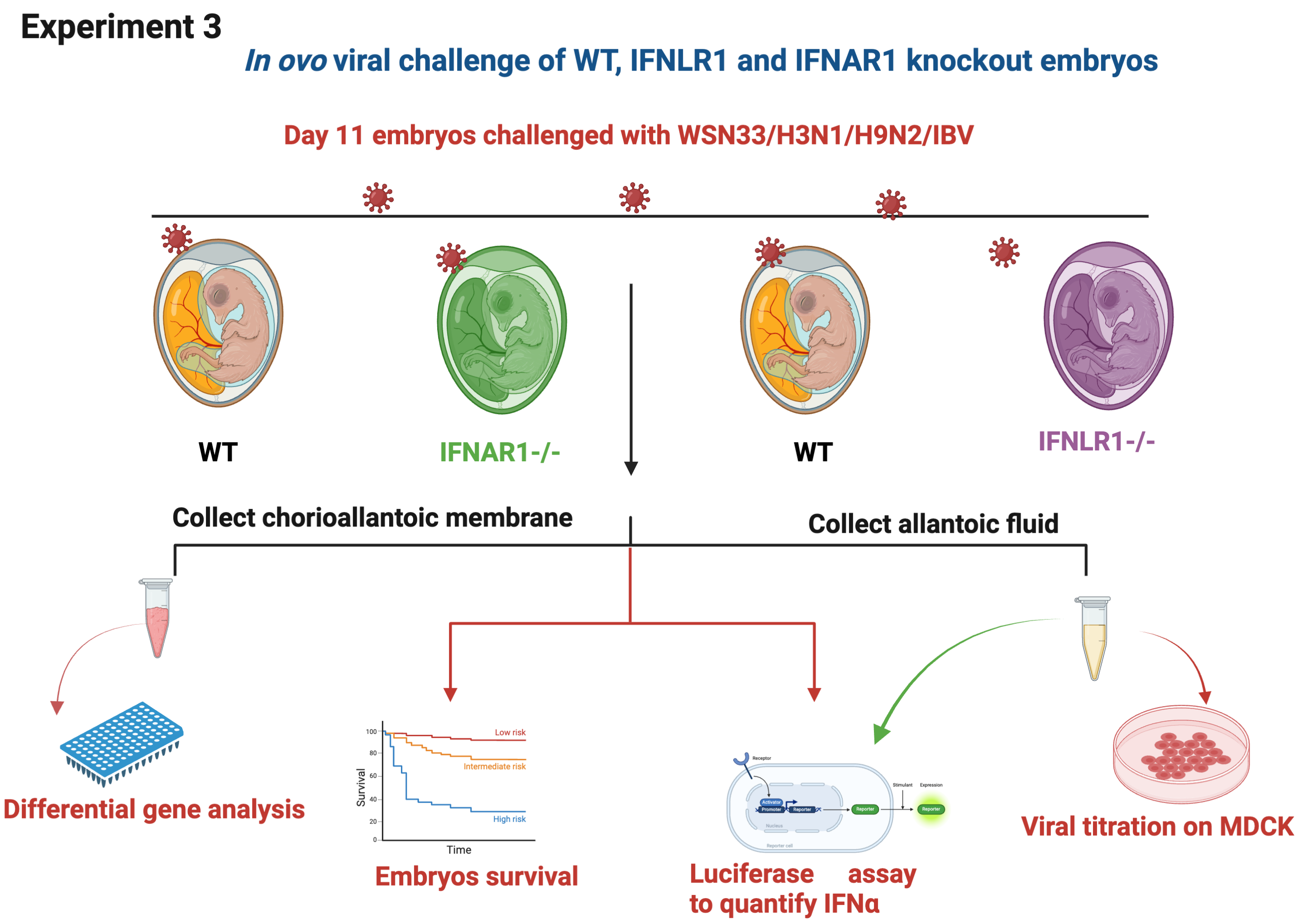
Experiment 3
*In ovo* viral challenge of WT, IFNLR1 and IFNAR1 knockout embryos
Day 11 embryos challenged with WSN33/H3N1/H9N2/IBV
WT
IFNAR1-/-
WT
IFNLR1-/-
Collect chorioallantoic membrane
Collect allantoic fluid
Differential gene analysis
Low risk
Intermediate risk
High risk
Survival
Time
Embryos survival
Luciferase assay to quantify IFNα
Viral titration on MDCK

Supp Fig. 9

**Supplementary Fig. 9.** Design of experiment 3, 11-day-old embryos were infected with 1000 FFU of four distinct viral strains: a laboratory-adapted human influenza A virus (H1N1), two low pathogenic avian influenza A virus strains (H3N1 and H9N2), and infectious bronchitis virus (IBV)- Beaudette strain virus. Allantoic fluid and chorioallantois membrane (CAM) were sampled 24 hours post-infection. The experiment was performed blindly, and the genotype of embryos was determined, and only WT and homozygous embryos were selected for further analysis. The viral titers of WSN33, H3N1, and H9N2 were quantified by titration on MDCK cells. For the IBV, qPCR was used to measure the viral titer in the CAM. Relative mRNA expression of Mx and IFN-λ were studied in the CAM using qPCR. IFN-α/β concentration was quantified by titration of allantoic fluids on CEC-32 #511 cells. The probability of survival in the challenged groups was also investigated.

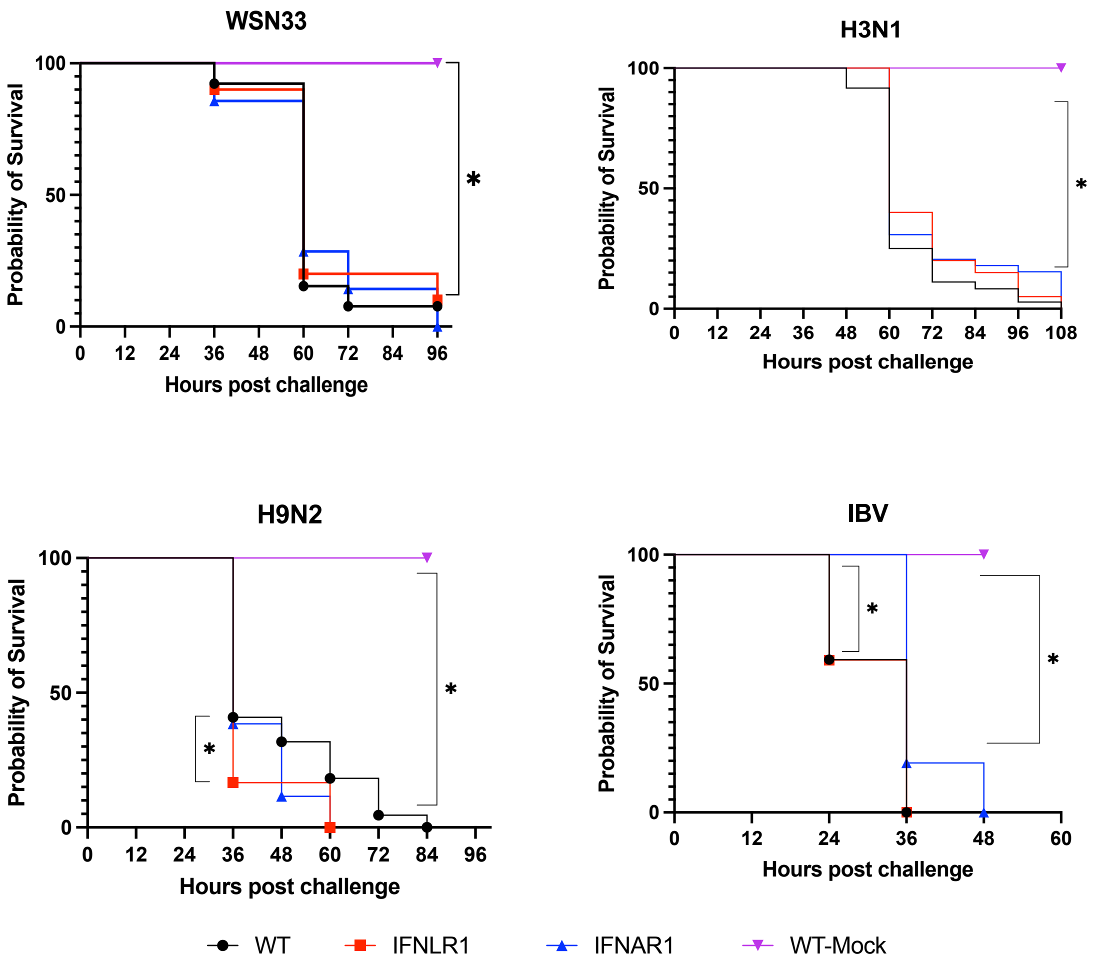
WSN33
H3N1
H9N2
IBV
Probability of Survival
Hours post challenge
100
50
0
0
12
24
36
48
60
72
84
96
108
*
WT
IFNLR1
IFNAR1
WT-Mock

**Supp Fig. 10**

**Supplementary Fig. 10**. The probability of survival in WT, IFNLR1$^{-/-}$ and IFNAR1$^{-/-}$ embryos. On embryonic day 11, chicken embryos were infected with 1000 FFU of influenza A virus strains (WSN33/H3N1/H9N2) and IBV- Beaudette strain, and the probability of survival in the challenged groups was recorded. Statistical differences between groups are indicated by asterisks (*****, $P \leq 0.05$).

#### Experiment 4

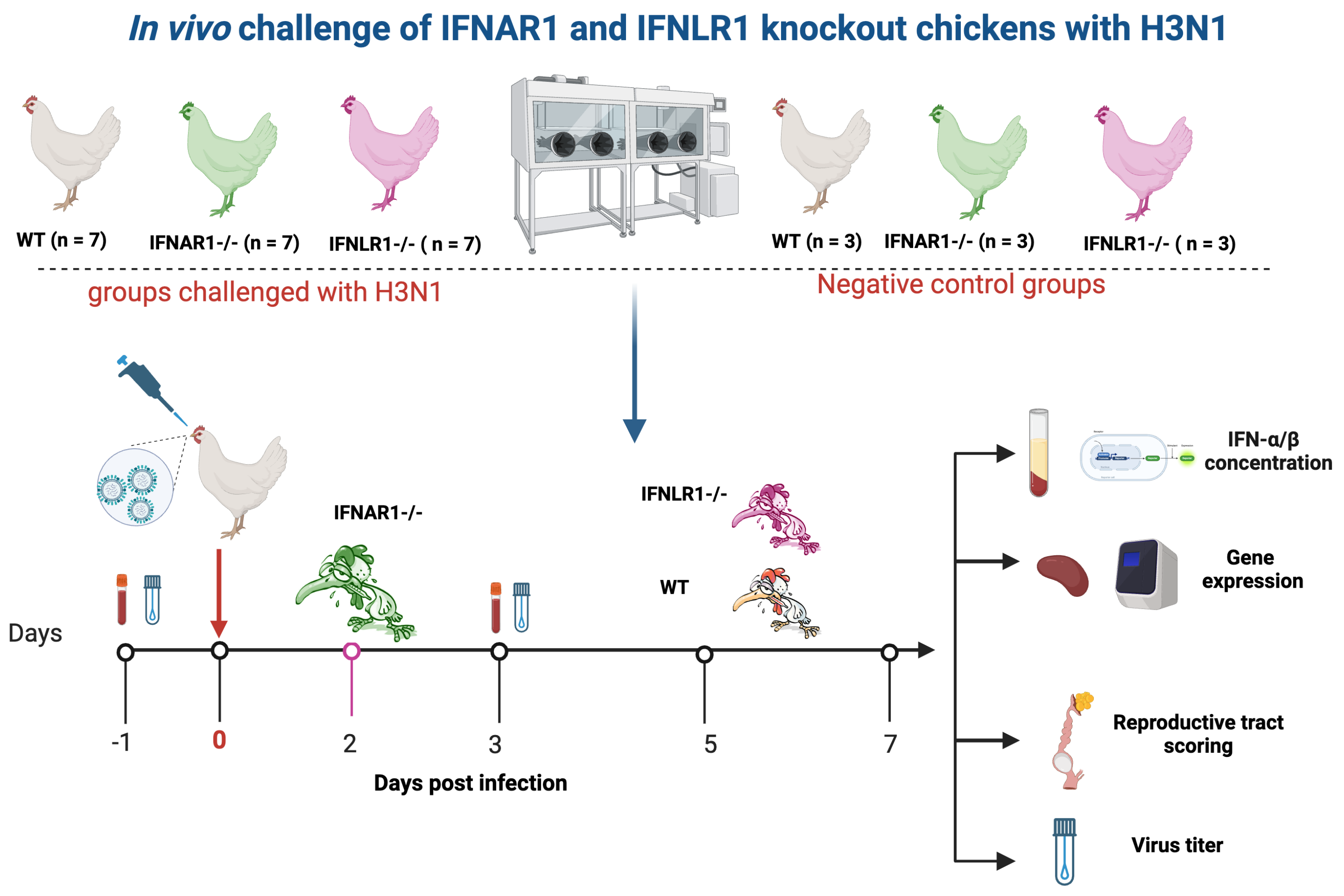


**Supp Fig. 11**

**Supplementary Fig. 11.** In experiment 4, 27-week-old WT, IFNLR1$^{-/-}$, and IFNAR1$^{-/-}$ hens were challenged with H3N1 avian influenza ($10^6$ FFU) in 0.2 ml PBS per bird and distributed via nasal and tracheal routes. Seven birds were used for the infection experiment, and three others were used as negative control per group. The IFNAR1$^{-/-}$ showed severe sickness symptoms and were euthanized at day 2 post-infection, whereas WT, IFNLR1$^{-/-}$ showed similar symptoms and were euthanized in the period between days 5-7 post-infection. Birds were monitored daily for clinical symptoms. Tracheal and cloacal swabs were collected to analyze the viral RNA loads by qPCR. Plasma was collected to estimate IFN-α/β concentration through luciferase assay. The spleen was collected for an immune-related gene expression study through qPCR. The female reproductive scoring was also studied to evaluate the impact of type I and type III IFNs on H3N1 pathogenesis.

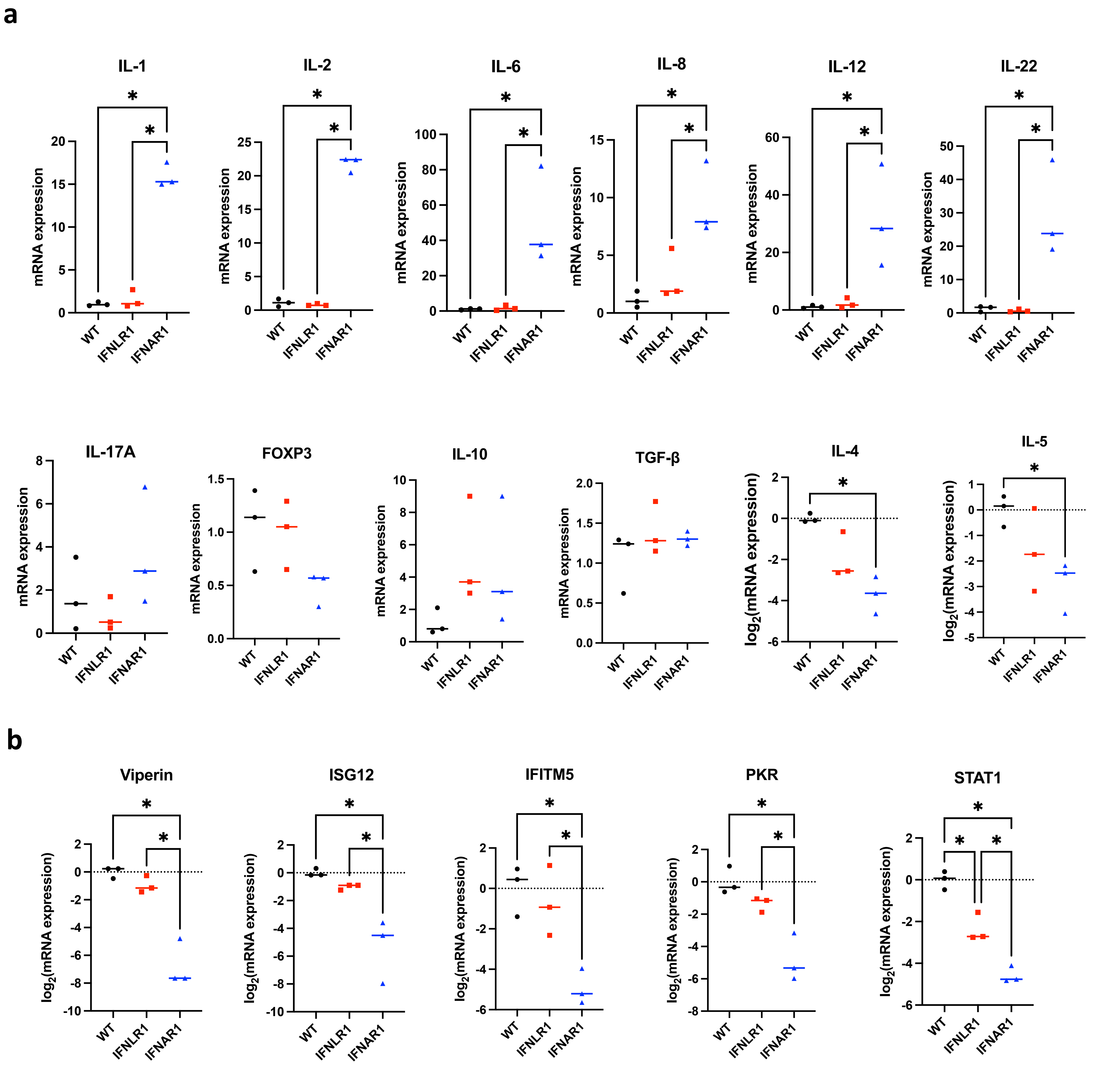
a
IL-1
IL-2
IL-6
IL-8
IL-12
IL-22
IL-17A
FOXP3
IL-10
TGF-β
IL-4
IL-5
mRNA expression
log2(mRNA expression)
WT
IFNLR1
IFNAR1
*
b
Viperin
ISG12
IFITM5
PKR
STAT1
log2(mRNA expression)
WT
IFNLR1
IFNAR1
*

c

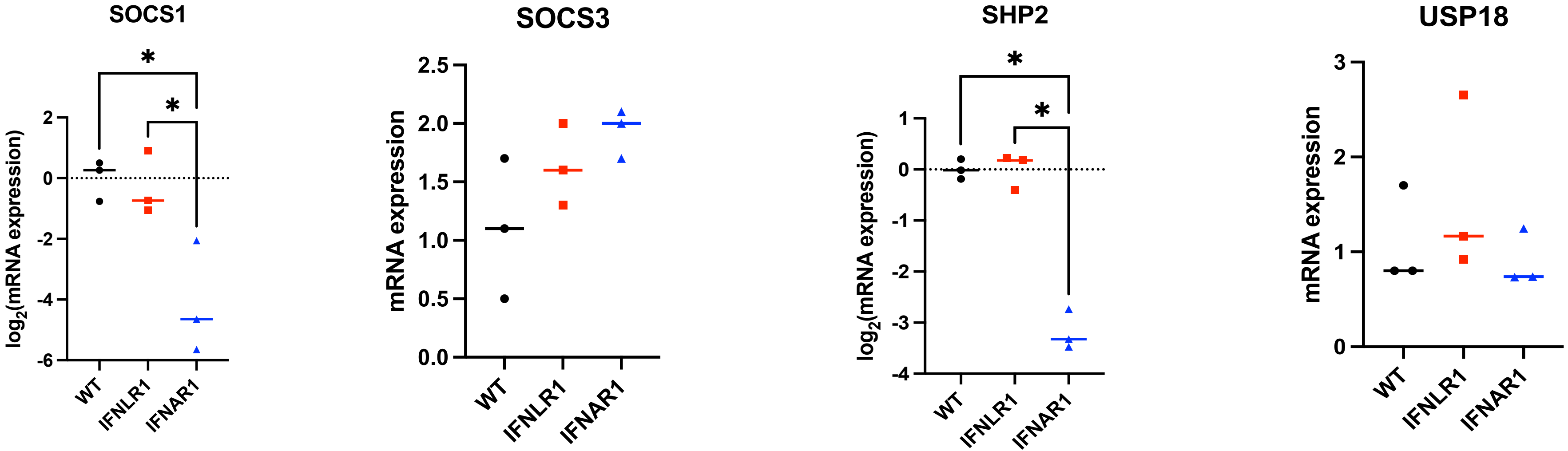

SOCS1
log2(mRNA expression)
2
0
-2
-4
-6
*
*
WT
IFNLR1
IFNAR1
SOCS3
mRNA expression
2.5
2.0
1.5
1.0
0.5
0.0
WT
IFNLR1
IFNAR1
SHP2
log2(mRNA expression)
1
0
-1
-2
-3
-4
*
*
WT
IFNLR1
IFNAR1
USP18
mRNA expression
3
2
1
0
WT
IFNLR1
IFNAR1


Supp Fig. 12

**Supplementary Fig. 12. a-c, Relative mRNA expression of various pro- and anti-inflammatory cytokines (a), interferon-stimulated genes (b), and interferon signaling inhibitors (c) in the spleen of H3N1 challenged hens**. For the qPCR study, RNA was isolated by Trizol and reverse transcripted with Go Script, and qPCR was done via Go Taq. The WT control group was used as a calibrator for the expression of all the studied genes, and the housekeeping gene 18S was used to normalize target gene expression. Relative quantification of the studied gene was performed using the $2^{\wedge\Delta\Delta CT}$ method. A no-template control was also included. The IFNAR1$^{-/-}$ hens were euthanized on day 2 post-infection. WT and IFNLR1$^{-/-}$ hens were euthanized on days 5 to 7 post-infection based on the days of euthanization. Only spleen samples collected on day 5 post-infection were included for WT and IFNLR1$^{-/-}$ hens for the gene expression study. Statistical differences between groups are indicated by asterisks (*****, $P \leq 0.05$).

**Supplementary Table 1.** List of designed sgRNA and ssODN

| Gene name | Type | Purpose | Sequence (5′ - 3′) |
|---|---|---|---|
| IFNLR1 sgRNA | single guide RNA | CRISPR/Cas9 targeting | GGCAGCCCAGAGGTACGTCA (PAM: TGG) |
| IFNAR1 sgRNA | single guide RNA | CRISPR/Cas9 targeting | GTGTGCCTCTGGGCGGCTAG (PAM: CGG) |
| IFNAR1 ssODN part 1 | repair construct | HDR | GCAGTCGTCAGAGGCTTCCGGTA AGGGTGAGCGCAGCCACGGACTG ATGGCTGAGGCGGCGTGTGCCTCT |
| IFNAR1 ssODN part 2 | repair construct | HDR | TAGCGGCTGTGCTGCTTTGTGTCCT GGTCGTGGTGTCCCGGTGCTGTGCA GGTGAGCCGAGCACGGTGCA |

**Supplementary Table 2.** Primers and probes were used for the genotype of IFNAR1 and IFNLR1 KO chicken.

| Genes | Internal name & Sequence (5′→3′) of Primers/Probes | Tm °C |
|---|---|---|
| IFNLR1 Primers | FW (1316): 5' GTTCTGCTTCAGCATGTCTGCAT 3'<br>RV (1317): 5' TCCAACCTCAGCCATACCACG 3' | 59 |
| IFNAR1 Primers | FW (1263): 5'- CGCAGCCACGGACTGAT -3'<br>RV (1264): 5'- ACGACCAGGACACAAAGCA-3' | 60 |
| IFNAR1 Probes | IFNAR1 MUT-probe_HEX (1291):<br>[HEX]CAGCCGCTAAGAGGCA[MGBEQ]<br>IFNAR1 WT-probe_FAM (1262):<br>[FAM]CCGCTAGCCGCCCAG[MGBEQ] | 60 |

**Supplementary Table 3.** Primers used for sexing and various RT-PCR assays and IBV qPCR.

| Purpose | Internal name & Sequence of Primers | Tm °C |
|---|---|---|
| Sexing | Z chromosome FW (1): 5'- AAGCATAGAACAATGTGGGAC-3'<br>Z chromosome RV (2): 5'- AACTCTGTCTGGAAGGACTT-3'<br>W chromosome FW (3): 5'- CTATGCCTACCACMTTCTATTTGC3'<br>W chromosome RV (4): 5'- AGCTGGAYTTCAGWCATCTTCT-3' | 56 |
| β-actin RTPCR | FW (277): 5'- TACCACAATGTACCCTGGC-3'<br>RV (278): 5'- CTCGTCTTGTTTTATGCGC-3' | 56 |
| IL-28Rα RTPCR | FW (1642):5'AGTGCTGGCAACTCTGTGCT3'<br>RV (1643): 5'TCCTTCTCCTGGAGTCCATGTCA3' | 62 |
| IBV qPCR | FW (1141): 5- GCTTTT GAGCCTAGC GTT-3'<br>RV (1142): 5'-GCCATGTTG TCACTG TCTATTG-3'<br>IBV probe (1140): 5'-FAM-CACCACCAGAACCTGTCACCTC-BHQ1-3' | 59 |
| 28S qPCR | FW (1277): 5- GGCGAAGCCAGAGGAAACT -3'<br>RV (1278): 5'-GACGACCGATTTGCACGTC -3'<br>28S probe (1279): 5'-HEX-AGGACCGCTACGGACCTCCACCA-BHQ1-3' | 59 |

**Supplementary Table 4.** Virus strains used for the *in ovo* challenge.

| Virus strain | Full name | Titer (U/µl) | Infection dose (U/100µl) |
|---|---|---|---|
| WSN | WSN/33 (H1N1) | 4500 | 1000 |
| H3N1 | A/Chicken/Belgium/460/2019 | 21000 | 1000 |
| H9N2 | A/chicken/Saudi Arabia/CP7/1998 | 24000 | 1000 |
| IBV | Infectious bronchitis Virus Beaudette strain | 1400 | 1000 |

**Supplementary Table 5.** clones and concentrations of antibodies used for the FACS studies of experiment 1 and experiment 2.

| **Primary Antibody** | **Internal code** | **Company** | **Clone** | **Concentration** |
|---|---|---|---|---|
| Experiment 1: FACS analysis of B cells, monocytes, γδ T cells (TCR1), αβ T cells (TCR2 + TCR3) and CD4-CD8 subsets of T cells | | | | |
| mouse IgG1 Anti-chicken TCRγδ-BIOT | 42 | Biozol | TCR-1 | 0.625 µg/mL |
| mouse IgG1 anti-chicken TCRαβ/Vb1-BIOT | 43 | Biozol | TCR-2 | 2.5µg/mL |
| mouse IgG1 anti-chicken TCRαβ/Vb2-BIOT | 44 | Biozol | TCR-3 | 2.5µg/mL |
| mouse IgG1 anti-chicken Bu1-FITC | 25 | Biozol | AV20 | 2.5µg/mL |
| mouse IgG1 anti-chicken KUL01_UNLAB | 23 | Biozol | KUL01 | 2.5µg/mL |
| mouse-IgG1_anti-chicken CD8α_PacBlue | 60 | Biozol | CT-8 | 0.625 µg/mL |
| mouse IgG2a anti-chicken CD8β_UNLAB | 61 | Biozol | EP42 | 0.625 µg/mL |
| mouse IgG1 anti-chicken CD4_FITC | 59 | Biozol | CT-4 | 0.625 µg/mL |
| **Secondary Antibody** | | **Company** | **Clone** | **Concentration** |
| rat anti-mouse IgG2a_PE | 62 | Biozol | SB84a | 0.125 µg/mL |
| Streptavidin_APC | 45 | VWR | - | 0.2 µg/mL |
| goat anti-mouse IgG (H+L)-APC | 26 | Biozol | polyclonal | 0.625 µg/mL |
| Fixable Viability Dye | 13 | eBioscience | ---- | 1:1000 |

| **Primary Antibody** | **Internal code** | **Company** | **Clone** | **Concentration** |
|---|---|---|---|---|
| eFluor 780 | | | | |
| Experiment 2: FACS analysis of B cells, monocytes, γδ T cells (TCR1), αβ T cells (TCR2 + TCR3), MHCI, MHCII, MHCII+ B cells and MHCII+ monocytes | | | | |
| mouse IgG1 anti-chicken Bu-1-AF647 | 22 | Biozol | AV20 | 1:500 |
| mouse IgG1 anti-chicken MHC I | 123 | southern Biotech | F21-2 | 1:200 |
| mouse IgG1 anti-chicken MHC II-AF488 | 118 | Biozol | 2G11 | 1:1000 |
| The primary antibodies utilized in both experiments were 42 (TCR1), 43 (TCR2), 44 (TCR3), and 23 (KUL01), alongside secondary antibodies 26 and 45, as well as the live-dead antibody 13. | | | | |

**Supplementary Table 6.** Details of various primers used for the qPCR study. The annealing temperature for all the used primers is 59 °C.

| Primer | Sequence 5' to 3' | amplicon (bp) | Source | Accession number |
|---|---|---|---|---|
| FoxP3 FW | AGTACGCCACAACCTGAGCCT | 157 | 1 | MT133687.1 |
| FoxP3 RV | TTGGGGTCCTCTCAGCTCCGT | | | |
| IL-12p35 FW | TGGCCGCTGCAAACG | 240 | 2 | NM_001398447.1 |
| IL-12p35 RV | CCAGCTCTGCCTTGTAGGTT | | 3 | |
| IL-17A FW | TTTCTGCACATGGGAAGGTG | 144 | 1 | AJ493595 |
| IL-17A RV | CCTGGTTCATGTTGCTGATGC | | | |
| IL-2 FW | GAACCTCAAGAGTCTTACGGGTCTA | 111 | 4 | NM_204153.2 |
| IL-2 RV | ACAAAGTTGGTCAGTTCATGGAGA | | | |
| IL-22 FW | TGTTGTTGCTGTTTCCCTCTTC | 143 | 1 | NM_001199614.1 |
| IL-22 RV | GCCAAGGTGTAGGTGCGATTCC | | | |
| IL-4 FW | GTGCCCACGCTGTGCTTAC | 82 | 1 | AJ621249. |
| IL-4 RV | AGGAAACCTCTCCCTGGATGTC | | | |
| IL-5 FW | GGAACGGCACTGTTGAAAAATAA | 111 | 1 | NM_001007084.2 |
| IL-5 RV | TTCTCCCTCTCCTGTCAGTTGTG | | | |
| IL-6 FW | GCTTCGACGAGGAGAAATGC | 139 | 1 | NM_204628 |
| IL-6 RV | GCCAGGTGCTTTGTGCTGTA | | | |
| TGF-β FW | CGGCCGACGATGAGTGGCTC | 120 | 1 | M31160.1 |
| TGF-β RV | CGGGGCCCATCTCACAGGGA | | | |
| IL-10 FW | CGGGAGCTGAGGGTGAAGT | 88 | 5 | AJ621254.1 |
| IL-10 RV | GTTCAGAGCTGAGCAGTTGGATGT | | | |
| IL-1β FW | GTGAGGCTCAACATTGCGCTGTA | 214 | 1 | NM_204524.2 |
| IL-1β RV | TGTCCAGGCGGTAGAAGATGAAG | | | |
| Viperin FW | CAGTGGTGCCGAGATTATGC | 105 | 6 | EU427332.1 |
| Viperin RV | CACAGGATTGAGTGCCTTGA | | | |
| ISG12 FW | TCCTCAGCCATGAATCCGAACA | 114 | 6 | BN000222.1 |

| | | | | |
|---|---|---|---|---|
| ISG12 RV | GGCAGCCGTGAAGCCCAT | | | |
| Mx FW | GCTCCTTCAGGAACTTCCGCTT | 115 | 1 | NM_204609.2 |
| Mx RV | TTCCCAGAGTTCCGGTCTCCAA | | | |
| IFN-λ FW | CTTTGGAGTTGAAGGCAGTGTGG | 195 | 7 | XM_040703743.1 |
| IFN-λ RV | TCTGGGTTGTGGGGTTTGTGAG | | | |
| STAT1 FW | TTGTAACTTCGCTATTGGTATTCC | 106 | 8 | NM_001012914 |
| STAT1 RV | TTCCGTGATGTGTCTTCCTTC | | | |
| TLR3 FW | TCAGTACATTTGTAACACCCCGCC | 256 | 9 | NM_001011691 |
| TLR3 RV | GGCGTCATAATCAAACACTCC | | | |
| MDA5 FW | CTCTGCGAGAAACCCAACAT | 329 | 10 | GU570144 |
| MDA5 RV | GCCCTCTGCTTCATCTTCAC | | | |
| MyD88 FW | ATCCCTCATTTCTGGCATCTT | 89 | 11 | AJ851640.1 |
| MyD88 RV | CCTTCCTTATAGTTCTGGCTTCT | | | |
| IFITM5 FW | TGCTTCACCAGCTAGGACTCTGC | 140 | 12 | XM_421662.4 |
| IFITM5 RV | TGGCTTTTGCTCTGTCACCACTTTG | | | |
| IL8 FW | TTGGAAGCCACTTCAGTCAGAC | 120 | 13 | NM_205498 |
| IL8 RV | GGAGCAGGAGGAATTACCAGTT | | | |
| IRF7 FW | CAGGAAGGATGTCACCAGCA | 117 | 14 | NM_205372.1 |
| IRF7 RV | GCGCAGCGGAAGTTGGTCTT | | | |
| NF-kB1 FW- | GGACGGCGAAAGGACTCT | 208 | 15 | NM_205134.2 |
| NF-kB1 RV- | CCATTGCAAACATTTGGGGAT | | | |
| SOCS3 FW | GCACCAAGAACCTGCGCATC | 103 | This study | NM_204600.2 |
| SOCS3 RV | AGCTTCAGCACGCAGTCGAA | | | |
| USP18 FW | GAGCACCTGGCCTGTCAGAT | 127 | This study | XM_040658339.2 |
| USP18 RV | AGCACTGCAGGCACTTCTCC | | | |
| SHP2 FW | TTGCAACTCAAGCAGCCCCT | 119 | This study | NM_204968 |
| SHP2 RV | TCCCAGAAGCCCTGCTTGAC | | | |
| SOCS1 FW | AGGGTTTTGCAGCCCTCGTT | 92 | This study | *NM_001137648.1* |

| | | | | |
|---|---|---|---|---|
| SOCS1 RV | GGCTCCCGTCCGTGCTAATT | | | |
| r18S FW | CATGTCTAAGTACACACGGGCGGTA | 136 | 16 | NC_052547. 1 |
| r18S RV | GGCGCTGCTGGCATGTATTA | | | |

**References:**


1. von Heyl, T. *et al.* Loss of αβ but not γδ T cells in chickens causes a severe phenotype. *European Journal of Immunology* **53**, 2350503 (2023).
2. Balu, S., Rothwell, L. & Kaiser, P. Production and characterisation of monoclonal antibodies specific for chicken interleukin-12. *Veterinary immunology and immunopathology* **140**, 140-146 (2011).
3. Abdul-Careem, M. *et al.* Marek's Disease Virus–Induced Transient Paralysis Is Associated with Cytokine Gene Expression in the Nervous System. *Viral immunology* **19**, 167-176 (2006).
4. Xu, F., Liu, S. & Li, S. Effects of selenium and cadmium on changes in the gene expression of immune cytokines in chicken splenic lymphocytes. *Biological Trace Element Research* **165**, 214-221 (2015).
5. He, S. *et al.* High-frequency and activation of CD4+ CD25+ T cells maintain persistent immunotolerance induced by congenital ALV-J infection. *Veterinary Research* **52**, 1-15 (2021).
6. Wang, S. *et al.* Dynamic changes in the expression of interferon-stimulated genes in joints of SPF chickens infected with avian reovirus. *Frontiers in Veterinary Science* **8**, 618124 (2021).
7. Yu, Y. *et al.* Effects of infectious bursal disease virus infection on interferon and antiviral gene expression in layer chicken bursa. *Microbial pathogenesis* **144**, 104182 (2020).
8. Truong, A.D., Hong, Y., Hoang, C.T., Lee, J. & Hong, Y.H. Chicken IL-26 regulates immune responses through the JAK/STAT and NF-κB signaling pathways. *Developmental & Comparative Immunology* **73**, 10-20 (2017).
9. Villanueva, A., Kulkarni, R. & Sharif, S. Synthetic double-stranded RNA oligonucleotides are immunostimulatory for chicken spleen cells. *Developmental & Comparative Immunology* **35**, 28-34 (2011).
10. Lee, S.B. *et al.* Targeted knockout of MDA5 and TLR3 in the DF-1 chicken fibroblast cell line impairs innate immune response against RNA ligands. *Frontiers in Immunology* **11**, 678 (2020).
11. Wang, Y. *et al.* Chicken interferon regulatory factor 7 (IRF7) can control ALV-J virus infection by triggering type I interferon production through affecting genes related with innate immune signaling pathway. *Developmental & Comparative Immunology* **119**, 104026 (2021).
12. Giotis, E. *et al.* Constitutively elevated levels of SOCS1 suppress innate responses in DF-1 immortalised chicken fibroblast cells. *Scientific Reports* **7**, 17485 (2017).
13. Li, Y., Handberg, K., Juul-Madsen, H.R., Zhang, M. & Jørgensen, P.H. Transcriptional profiles of chicken embryo cell cultures following infection with infectious bursal disease virus. *Archives of virology* **152**, 463-478 (2007).
14. Wu, W.-J. *et al.* Effects of Reticuloendotheliosis virus on TLR-3/IFN-B pathway in specific pathogen-free chickens. *Research in Veterinary Science* **156**, 36-44 (2023).
15. Kuehu, D.L. *et al.* Effects of Heat-Induced Oxidative Stress and Astaxanthin on the NF-kB, NFE2L2 and PPARα Transcription Factors and Cytoprotective Capacity in the Thymus of Broilers. *Current Issues in Molecular Biology* **46**, 9215-9233 (2024).
16. Laparidou, M., Schlickenrieder, A., Thoma, T., Lengyel, K. & Schusser, B. Blocking of the CXCR4-CXCL12 interaction inhibits the migration of chicken B cells into the bursa of Fabricius. *Frontiers in immunology* **10**, 3057 (2020).
